# Probabilistic Fine-mapping of Putative Causal Genes

**DOI:** 10.1101/2024.10.27.620482

**Authors:** Jeffrey Okamoto, Xianyong Yin, Brady Ryan, Joshua Chiou, Francesca Luca, Roger Pique-Regi, Hae Kyung Im, Jean Morrison, Charles Burant, Eric B. Fauman, Markku Laakso, Michael Boehnke, Xiaoquan Wen

## Abstract

Integrative genetic analysis of molecular and complex trait data, including colocalization analysis and transcriptome-wide association studies (TWAS), has shown promise in linking GWAS findings to putative causal genes (PCGs) underlying complex diseases. However, existing methods have notable limitations: TWAS tend to produce an excess of false-positive PCGs, while colocalization analysis often lacks sufficient statistical power, resulting in many false negatives. This paper introduces a probabilistic fine-mapping method, INTERFACE, which is designed to identify putative causal genes while accounting for direct variant-to-trait effects within genomic regions harboring multiple gene candidates. INTERFACE leverages interpretable, data-informed priors that incorporate both colocalization and TWAS evidence, enhancing the sensitivity and specificity of PCG inference and setting it apart from existing methods. Additionally, INTERFACE implements analytical measures to improve the accuracy of gene-to-trait effect estimation. We apply INTERFACE to METSIM plasma metabolite GWASs and UK Biobank pQTL data to identify causal genes regulating blood metabolite levels and demonstrate the unique biological insights INTERFACE provides.

## 1 Introduction

Genome-wide association studies (GWAS) have successfully identified an abundance of genetic variants underlying complex diseases. Building on these successes, integrative genetic association analysis has emerged as a powerful approach to exploring the molecular mechanisms underlying these diseases by examining shared genetic associations among molecular and complex traits. Common analytical tools for integrative genetic association analysis include transcriptome-wide association studies (TWAS) [1, 2, 3, 4], which assess the correlation between genetically-predicted gene expression and complex trait levels, and colocalization analysis, which identifies overlapping causal genetic variants underlying molecular and complex traits [5, 6, 7]. Both types of analysis have been extensively utilized to link GWAS findings to putative causal genes (PCGs). While TWAS can be applied to many types of omics data, including proteomic data [8, 9], we refer to this class of analyses as TWAS throughout the paper.

Despite their broad applications, both TWAS and colocalization analyses have noticeable limitations in identifying PCGs. TWAS analysis is connected to instrumental variable and Mendelian randomization analyses, and interpreting TWAS results as indicative of PCGs requires assumptions that are difficult to verify. In practice, many TWAS signals represent spurious PCGs due to patterns of linkage disequilibrium (LD), commonly referred to as indirect horizontal pleiotropy or LD hitchhiking [10, 11, 12]. Hukku et al. [11] demonstrate that LD between molecular QTLs and GWAS signals, rather than causal molecular mechanisms, often explains the majority of TWAS signals. While colocalization analysis does not rely on causal assumptions, its interpretation is also limited. For example, it cannot distinguish between horizontal and vertical pleiotropy [13]. Moreover, colocalization analysis often lacks power largely because genetic association analyses frequently struggle to pinpoint the causal variants of either molecular or complex traits due to limited sample sizes and complex LD patterns [14].

Recently, two strategies have been developed to enhance the effectiveness of PCG identification methods. The first jointly models the causal effects of multiple neighboring genes and SNPs within the TWAS framework [10, 12, 15]. The second explicitly combines colocalization and TWAS evidence [5, 11, 16]. Both strategies have shown promise in mitigating LD hitchhiking effects and improving the specificity of PCG identification. However, there is still considerable room for improvement. The methods using joint modeling often encounter inferential challenges due to the identifiability of the underlying statistical model and computational complexity, requiring unrealistic simplifying assumptions to address these obstacles. Existing methods that combine colocalization and TWAS evidence typically analyze one gene at a time. As a result, they are suboptimal to effectively handle cases where multiple genes exhibit highly correlated molecular phenotypes and fail to maximize the power for identifying PCGs.

In this paper, we introduce a computational method called INTERFACE (INTEgRative Fine mApping of Causal gEnes), which combines the strengths of the two aforementioned strategies for PCG discovery. INTERFACE performs probabilistic multi-gene fine-mapping analysis using a hierarchical model with priors informed by both TWAS and colocalization evidence. INTERFACE is built on a principled empirical Bayes framework and is computationally efficient. Additionally, we explore and implement analytical measures in INTERFACE to improve the estimation accuracy for gene-to-trait effects. We systematically evaluate INTERFACE’s performance through extensive simulation studies. Finally, we apply INTERFACE to UK Biobank plasma protein QTL data (*n* = 35, 000) and METSIM plasma metabolite GWAS data (*n* = 10, 188) to infer causal genes underlying plasma metabolite levels. Our software implementation of INTERFACE is freely available at https://github.com/jokamoto97/INTERFACE/.

## 2 Results

### 2.1 Method Overview

INTERFACE implements a Bayesian variable selection algorithm that simultaneously considers multiple candidate causal genes and direct-effect causal genetic variants in a genomic region. The relationship between INTERFACE and existing single-gene PCG implication methods (e.g., TWAS and INTACT) is analogous to the relationship between multi-SNP fine-mapping methods and single-variant analyses in genetic association analysis. Similar to multi-SNP fine-mapping methods, INTERFACE improves the sensitivity for detecting both gene- and variant-level associations while accounting for complex LD structures. Additionally, INTERFACE uniquely quantifies probabilistic evidence from observed data across a comprehensive set of possible scenarios, where multiple genes and/or genetic variants may independently influence the complex trait of interest.

The INTERFACE method is based on a structural equation model (SEM), generalizing many existing TWAS-based methods, including INTACT [16], FUSION [1], and cTWAS [12]. Consider a genomic region harboring *p* genetic variants and *q* candidate genes. Let ***G*** (*n* × *p*), ***M*** *_i_* (*n* × 1), and ***Y*** (*n* × 1) denote the genotype matrix for the *p* variants, the molecular trait levels (e.g., expression levels or protein abundance) of the *i*th gene, and the complex trait vector for *n* unrelated population samples, respectively. We use the following SEM, consisting of *q* + 1 linear models, to jointly model potential causal relationships from all *q* candidate genes and *p* genetic variants, i.e.,

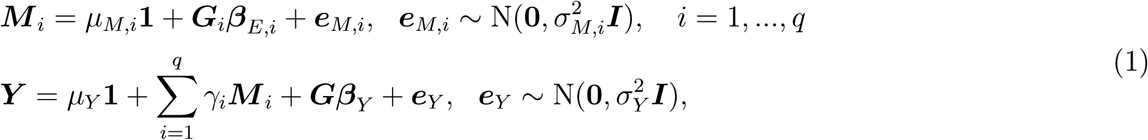

where ***G****_i_* (*n* × *p_i_*) denotes the *cis*-genotype matrix for the *i*th gene, ***β****_E,i_* denotes the *cis*-QTL effects vector for the *i*th gene, and the *p*-vector ***β****_Y_* denotes the direct (variant-to-trait) effects. The gene-to-trait effects, *γ_i_, i* = 1*, …, q*, are of primary interest for inference.

SEM (1) is closely related to the instrumental variable (IV) analysis/Mendelian randomization model that is widely applied in causal inference. One of the key advantages of this class of causal inference models is their ability to account for potentially unobserved confounding factors shared between molecular and complex traits through the residual errors (i.e., ***e****_Y_* and ***e****_M,i_*). Proper inference of gene-to-trait effects ***γ*** = (*γ*_1_ *…, γ_q_*) requires fitting a linear model where the molecular phenotype ***M*** *_i_* is replaced by their corresponding genetic predictions, ***M̂*** *_i_* = ***G****_i_****β̂****_E,i_*, for all genes, i.e.,

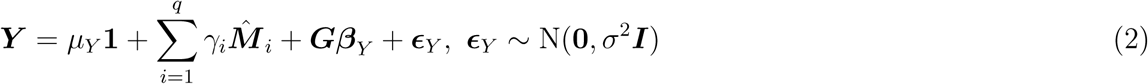

The causal interpretation of the inference results obtained by fitting Model (2) applies to both one-sample (i.e., molecular and complex traits are measured from the same set of samples) and two-sample (i.e., molecular and complex traits are measured from different sets of samples) designs under the usual causal assumptions of the IV analysis, including the inclusion, randomization, and exclusion assumptions. Furthermore, INTERFACE relaxes the exclusion assumption by explicitly modeling and controlling for the direct variant-to-trait effects (***β****_Y_*).

Performing inference of ***γ*** based on Model (2) presents significant challenges. Because the predicted molecular phenotypes ***M̂*** *_i_, i* = 1*, …, q* are functions of ***G***, the parameters ***γ*** and ***β****_Y_* can become non-identifiable [10, 16, 17]. Additionally, variable selection procedures require careful consideration, as the predictors in ***G*** and ***M̂*** *_i_* can be highly correlated. INTERFACE addresses these challenges by adopting an empirical Bayes framework. Most importantly, it pre-computes a set of interpretable informative priors for ***γ*** and ***β****_Y_* to resolve the identifiability issue. Moreover, INTERFACE utilizes the state-of-the-art variational inference algorithm, SuSiE [18], for Bayesian variable selection.

One of our main contributions is the development of interpretable informative priors for probabilistic causal gene fine mapping, integrating marginal TWAS and colocalization evidence. The priors for PCGs are based on the previous findings that colocalization and TWAS evidence are both *necessary* conditions for identifying genuine PCGs under the assumed SEM (1). The two sources of evidence are also considered practically Dempster-Shafer (D-S) independent, as genes with strong TWAS evidence often have little overlap with those showing strong colocalization evidence [11, 16]. Additionally, the priors for direct-effect SNPs can be derived from estimated molecular QTL enrichment information in colocalization analysis. In summary, all the required priors in INTERFACE are data-driven and well-justified. We provide the mathematical details of prior construction in the Methods section.

There has been limited effort in the literature to ensure the accurate estimation of gene-to-trait effects for PCGs. Our findings indicate that the quality of the estimates is heavily dependent on the strength of the genetic instruments used to predict molecular phenotypes. Specifically, including weak genetic instruments in the computation of ***M̂*** *_i_* can lead to poor estimation of gene-to-trait effects for gene *i*. These findings are consistent with similar observations in Mendelian randomization/IV analysis. Here, we propose using cumulative variant-level posterior inclusion probabilities of an independent signal cluster (or credible set) to gauge the instrumental strength of a corresponding molecular QTL. By implementing such a strategy to screen and select strong genetic instruments, INTERFACE effectively improves the accuracy of gene-to-trait estimates.

We summarize the INTERFACE workflow in Figure 1. Briefly, provided QTL and GWAS data, we perform marginal TWAS and colocalization analysis. We screen QTL fine-mapping results to remove weak genetic instruments from molecular trait prediction models. Gene prior inclusion probabilities are derived using the molecular QTL enrichment parameter estimates, gene-level colocalization probabilities from colocalization analysis, and the TWAS prior estimate. An exchangeable prior inclusion probability for direct-effect SNPs and the null model prior probability are also derived from molecular QTL enrichment estimates. Finally, we apply the variational inference algorithm from SuSiE [18] to fit the regression model (2), which includes all genetic variants and predicted molecular trait levels for all genes within the genomic region of interest as predictors. The output includes the posterior inclusion probabilities (PIPs), posterior mean effect sizes, and corresponding posterior standard deviations for each candidate gene.

**Figure 1:**
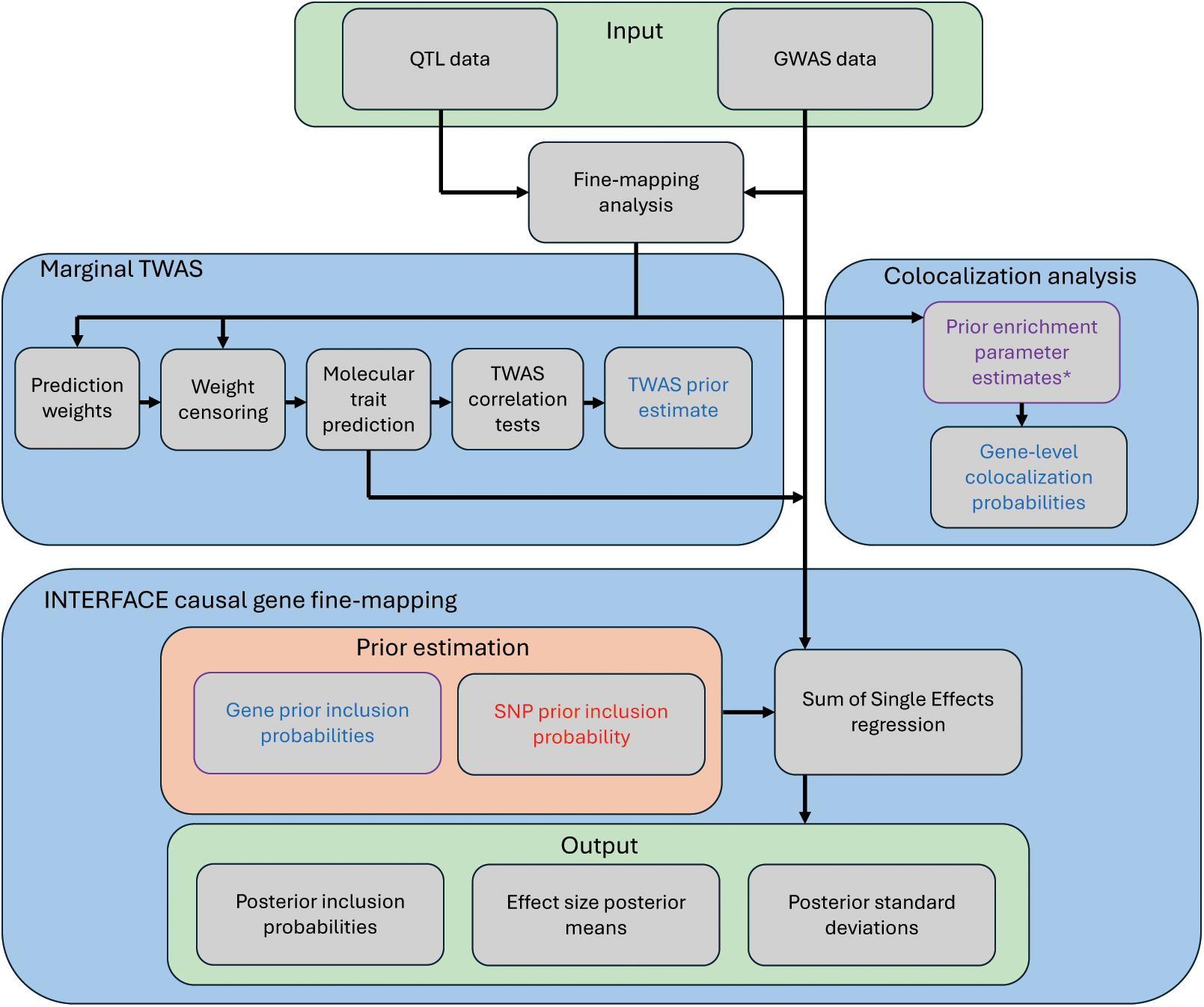
INTERFACE workflow. Text color indicates the quantities used for estimating each prior. Blue text denotes quantities used to estimate the gene prior inclusion probabilities (TWAS prior estimate and gene-level colocalization probabilities). Red text denotes quantities used to estimate the SNP prior inclusion probability. (*) Purple text denotes quantities used to estimate both gene and SNP prior inclusion probabilities (prior enrichment parameter estimates).

### 2.2 INTERFACE improves specificity and sensitivity in PCG identification

The SEM assumed by INTERFACE accommodates a range of possible causal scenarios, allowing for various combinations of variant-to-trait and gene-to-trait effects. This comprehensive approach enhances INTERFACE’s specificity and sensitivity in identifying PCGs over existing methods. Below, we present some practical scenarios for illustration.

One of the primary challenges in PCG identification is to distinguish between horizontal pleiotropy, where QTLs exert direct effects on complex traits, and vertical pleiotropy, where the molecular phenotype of interest mediates QTL effects on complex traits. INTERFACE addresses this challenge using a model comparison approach, which evaluates the evidence for competing scenarios by combining likelihood and prior information derived from the data. In the vertical pleiotropy scenario, the likelihood for candidate gene *i* characterizes the strength of the correlation, cor(***Y***, ***M̂*** *_i_*), while the corresponding prior accounts for the colocalization evidence for the target gene and the overall prevalence of vertical pleiotropy in the observed data. When there is only a single molecular QTL for the target gene, the likelihood for both vertical and horizontal pleiotropy scenarios are identical, and the prior breaks the tie. When multiple molecular QTLs/instruments of the target gene are available, the likelihood computation can provide crucial distinguishing information. Consider a hypothetical example in which a target gene contains two independent QTLs, but the two QTLs suggest opposite gene-to-trait effects. In such a case, cor(***Y***, ***M̂***) can be considerably weaker than the correlations obtained from the competing horizontal pleiotropic model. However, cor(***Y***, ***M̂***) can still be significantly different from 0, which may cause false positives for some standard TWAS methods. In this example, the INTERFACE model implicitly accounts for the heterogeneity of gene-to-trait effects by different instruments, akin to an automatic sensitivity diagnosis in instrumental variable analysis. We present a real data scenario that represents this hypothetical example in Section 2.4 and Figure 6. It is also worth noting that the probabilistic evidence reported by the INTERFACE posterior probabilities does not entirely dismiss one scenario over the other; rather, it quantifies their relative plausibility.

Jointly modeling multiple candidate genes offers several advantages for INTERFACE. First, when the region of interest contains multiple causal genes, fitting a joint model helps reduce residual error variance estimates and enhances the signal-to-noise ratio, thereby improving the sensitivity of identifying genuine PCGs. Second, it is often observed that multiple genes within a genomic region exhibit correlated expression levels and share molecular QTLs. In such cases, single-gene PCG implication methods often identify all involved genes as multiple independent PCGs. In contrast, INTERFACE considers and quantifies multiple plausible causal scenarios, including the possibility that the observed association pattern is driven by a single PCG, supported by its parsimony-inducing priors.

We conduct extensive simulations to evaluate the performance of INTERFACE using the real genotype data from the GTEx project [19]. The selected genomic region for simulation, previously studied by [11], has a complex LD structure (Supplemental Figure S1), making the simulated data susceptible to LD hitchhiking effects. The simulation scheme (detailed in the Methods section) is intentionally designed to differ from the INTERFACE model to test its robustness. The simulation results, summarized in Figure 2, indicate that INTERFACE achieves near-optimal power (compared to the oracle model) in identifying PCGs while effectively controlling the false discovery rate (FDR). In contrast, a class of TWAS methods generates excessive false positive findings, likely because they fail to account for direct variant-to-trait effects and horizontal pleiotropy. INTERFACE also outperforms the single-gene PCG implication methods, demonstrating much-improved power by simultaneously modeling multiple gene candidates. Furthermore, compared to alternative specifications of priors for Model (2) in a multi-gene fine-mapping setting, INTERFACE consistently shows superior performance (Panel B of Figure 2).

**Figure 2:**
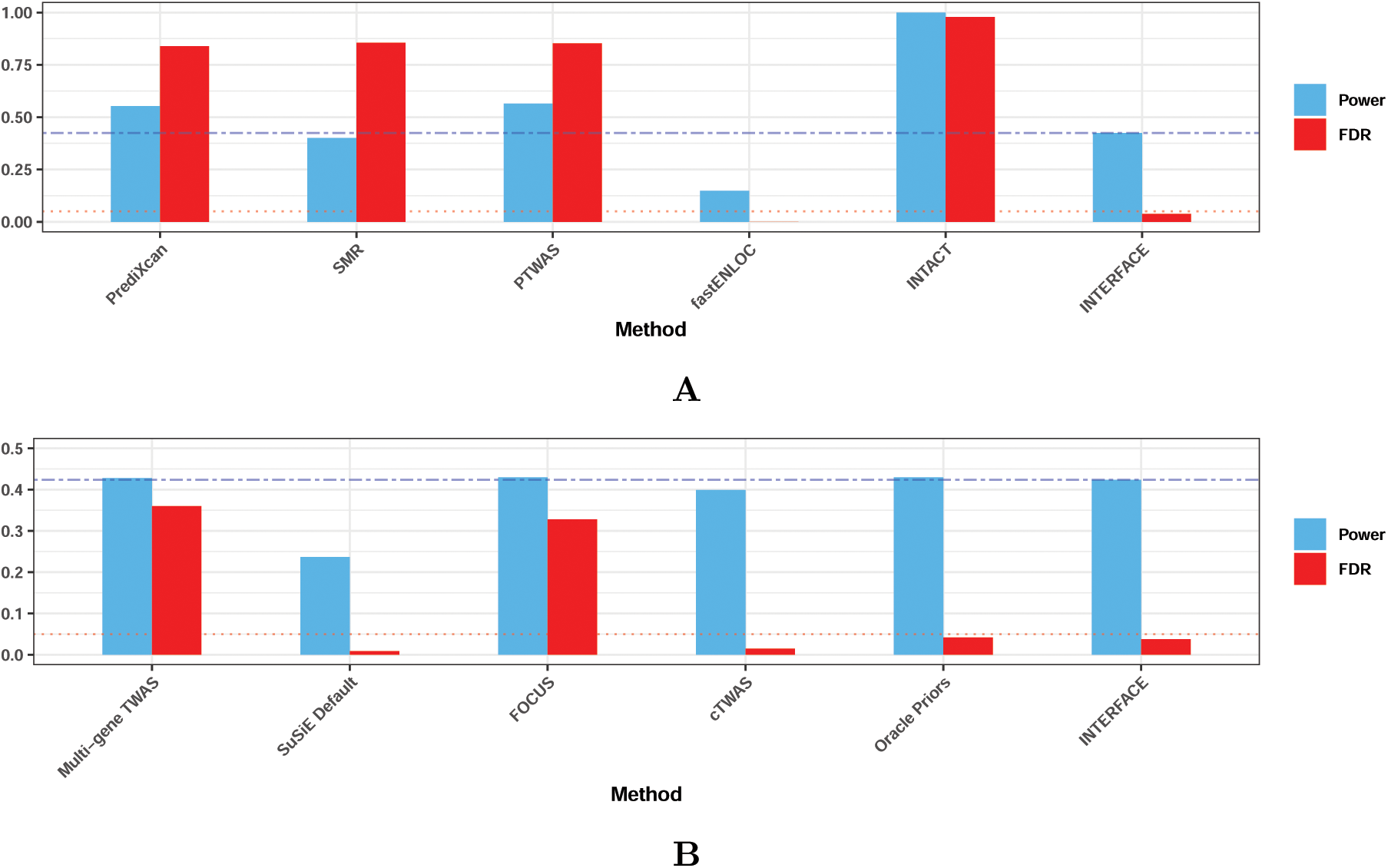
Comparison of FDR control and power in selecting putative causal genes. The realized false discovery rates (at the control level of 0.05) and the power to select PCGs by various methods are compared to INTERFACE using simulated data. In both panels, the red dotted line represents the target FDR control level, and the blue dashed line represents INTERFACE’s power. In the top panel, INTERFACE is compared to single-gene PCG implication methods; in the bottom panel, INTERFACE is compared to multi-gene methods with alternative prior specifications. Specifically, “Oracle priors” refer to the setting that uses the true prior probabilities for generating the simulated data.

### 2.3 INTERFACE accurately estimates gene-to-trait effects

We investigate INTERFACE’s estimation of gene-to-trait effects using simulated data, with a focus on the accuracy of the posterior expectation of ***γ*** by different genotype-based molecular phenotype prediction methods. Since INTERFACE does not impose restrictions on the form of ***M̂***, it allows for exploring various analytical approaches to construct predicted molecular phenotypes from genotype data. The default prediction models in our simulation studies are constructed by the PTWAS method [3].

Our key finding is that removing extremely weak instruments from the molecular phenotype prediction model can greatly improve the accuracy of gene-to-trait effect estimation, effectively eliminating outlying estimates assessed with large variance. Figures 3 and 4 indicate that extremely weak instruments are responsible for the outlying gene-to-trait effect estimates. Moreover, we identify that the cumulative PIP (CPIP) within a signal cluster representing the strength of evidence for an independent molecular QTL is an effective measure for gauging the strength of the corresponding genetic instrument. Based on these results, we implement the INTERFACE’s default effect estimation procedure by applying a threshold on CPIP values when constructing ***M̂***. Figure 3 illustrates the impacts of varying CPIP thresholds on effect estimate accuracy, power, and realized FDRs in selecting PCGs. Notably, including weak instruments does not inflate the realized FDR beyond its target control level, and thresholding CPIP only slightly reduces the power of PCG discovery. To achieve a balance between robust PCG identification and accurate estimation of gene-to-trait effects, we propose screening molecular QTLs for constructing ***M̂*** by a modest threshold. In INTERFACE’s default implementation, we use a CPIP threshold of 0.25.

**Figure 3:**
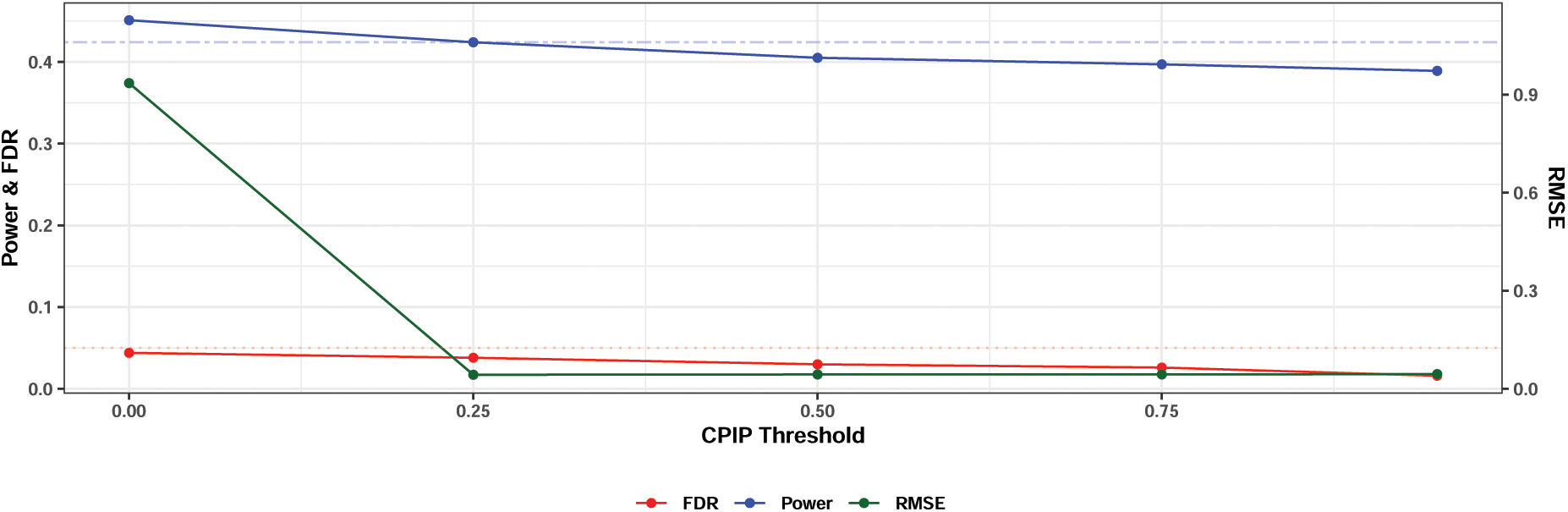
Impact of instrument selection on FDR, power, and accuracy of gene-to-trait estimates in INTERFACE. The dotted red line and the dashed blue line represent the target FDR control level and the reported INTERFACE power in the simulation study. Only SNPs within the signal clusters that pass the specified CPIP threshold are selected for predicting molecular phenotypes. Setting CPIP to 0 indicates no explicit selection procedure, while setting CPIP to 0.95 corresponds to the most stringent selection criterion. The figure shows that selecting genetic instruments has limited effects on the realized FDR and power; however, it is critical for ensuring the accuracy of estimating gene-to-trait effects.

**Figure 4:**
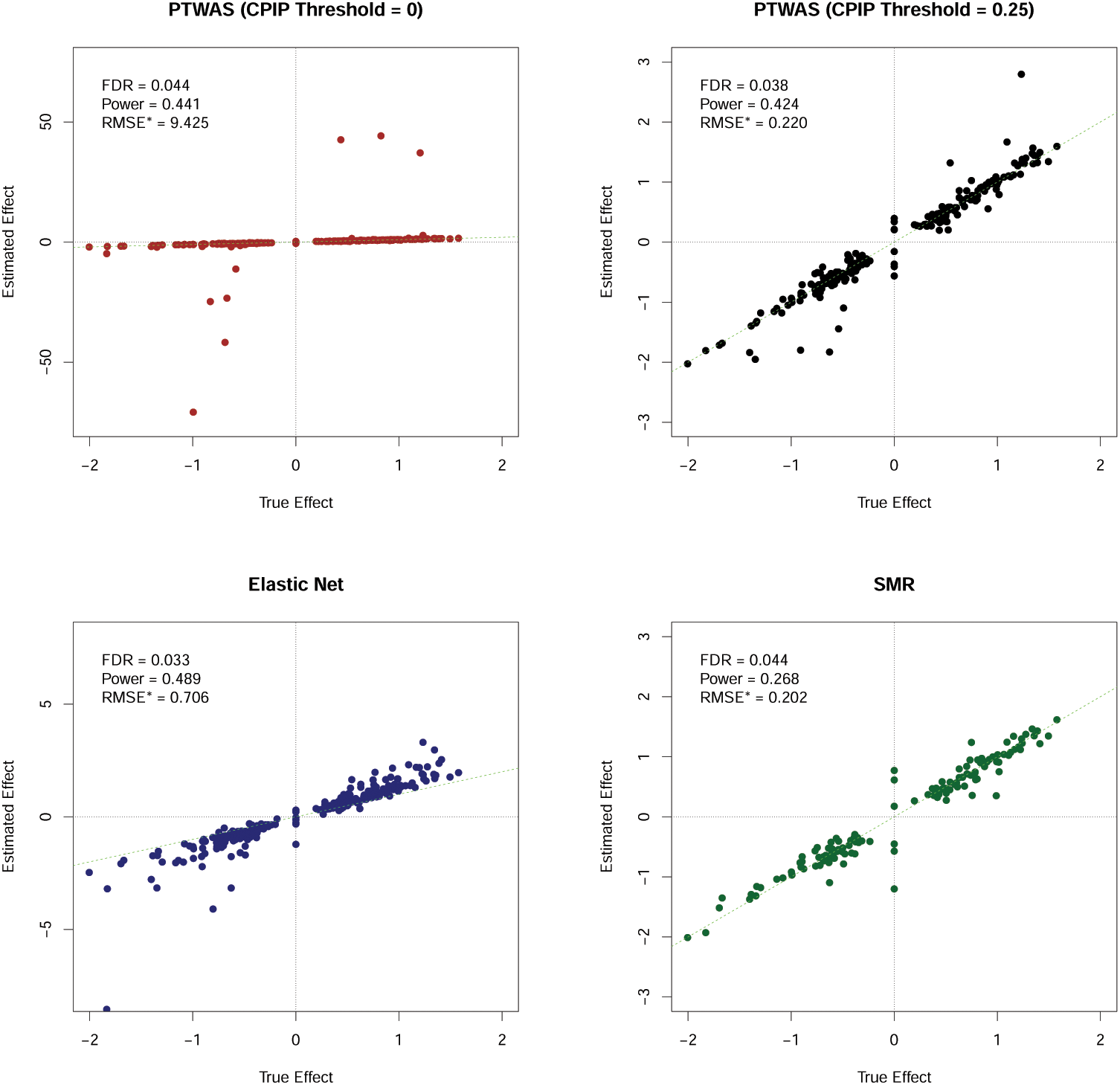
Accuracy of gene-to-trait effect estimation by INTERFACE with various molecular phenotype prediction models. The figure illustrates the effects of various genotype-based molecular phenotype prediction models on INTERFACE’s ability to identify potentially causal genes (PCGs) and the accuracy of the associated gene-to-trait effect estimates. Each prediction model provides INTERFACE with different molecular phenotype predictions and modifies the INTERFACE priors. The scatter plots show comparisons between the true gene-to-trait effects and the estimated values for genes that INTERFACE rejected at the FDR control level of 0.05. RMSE^∗^ denotes the root mean squared error of the estimated effects for the INTERFACE-rejected genes. The corresponding realized FDR and power are also labeled for each prediction method. In summary, INTERFACE properly controls FDR across all prediction methods but with varying power. SMR, relying on a single instrument for prediction, has the lowest power. The accuracy of the effect estimates also vary across methods (note the *y* scale difference in the 4 plots). Comparing PTWAS methods with CPIP thresholds of 0 and 0.25 (the top panel) highlights the importance of instrument selection for the accuracy of the gene-to-trait estimates.

In many practical cases, only 95% credible sets from fine-mapping analyses are made available as summary statistics. Figure 3 supports the use of these high-confidence association results for INTERFACE analysis, which should yield high-specificity PCGs and accurate gene-to-trait effect estimates, though with some potential loss of power.

Additionally, we examine alternative ***M̂*** for constructing methods, assessing their ability to identify PCGs and the accuracy of the corresponding gene-to-trait estimates (Figure 4). Prediction models that use shrinkage methods, such as the elastic-net method in PrediXcan, demonstrate improved power compared to the default PTWAS predictions. However, the resulting gene-to-trait estimates show systematic bias and have relatively large RMSE. Methods using only the strongest available instruments, such as SMR, perform well in estimating large effects but tend to underestimate small to modest effects, leading to much reduced power for identifying PCGs.

### 2.4 Analysis of METSIM Metabolon metabolite GWAS data

We apply INTACT and INTERFACE to jointly analyze the UK Biobank plasma pQTL data [20] and the Metabolic Syndrome in Men (METSIM) metabolite GWAS data [21], aiming to identify causal genes regulating human metabolite levels. The METSIM data comprises whole-genome sequencing data and measurements of 1,408 plasma metabolites from 10,188 Finnish men. For UK Biobank pQTL data, we use summary information in the form of 95% credible sets for each fine-mapped high-confidence pQTL. Prior to the integrative analysis with the pQTL data, single-SNP genetic association and fine-mapping analyses are performed on the metabolite data. Subsequently, we conduct colocalization and PWAS analyses based on the inferred 95% credible sets of metabolite GWAS hits and pQTLs using fastENLOC and PTWAS, respectively.

In the INTERFACE analysis, we first identify 193 genomic regions potentially harboring metabolite PCGs by applying INTACT at a relaxed FDR level. These genomic regions represent 951 unique genes across 168 unique metabolites or 5,509 unique metabolite-gene pairs. On average, each region is 14 Mb in length and contains 19 genes. We then apply INTERFACE to each region and identify PCGs at the FDR 5% level. Among the 5,509 candidate metabolite-gene pairs, INTERFACE identifies 132 metabolite-PCG pairs, substantially more than the 84 pairs identified by the single-gene INTACT analysis using the same FDR control threshold. We also find that INTERFACE identifies the majority (84%) of findings identified by INTACT.

We validate the PCG-metabolite pairs identified by INTERFACE and INTACT using a knowledge-based approach (KBA), which represents a collection of causal genes annotated for each metabolite [21, 22, 23, 24]. In brief, the KBA links a target metabolite to causal genes near its strong GWAS signals based on the known biochemistry of metabolic pathways. In total, 153 of the 5,509 gene-metabolite pairs analyzed by the integrative approaches were nominated by the KBA. Since not all KBA nominations rely on proteomic evidence, we expect that PCG findings from the integrative analysis of metabolite GWAS and pQTL data represent only a fraction of the KBA causal genes. In our analysis, INTACT identifies 53 (∼ 35%), and INTERFACE identifies 77 (∼ 50%) of these KBA findings. Considering that the KBA represents a high-quality set of true positive gene-metabolite pairs, we conclude that INTERFACE demonstrates improved sensitivity over INTACT for discovering PCGs in real data. The left panel of the Figure 5 illustrates the overlap of gene-metabolite pairs implicated by the KBA, INTACT, and INTERFACE. Importantly, we note that pairs implicated by INTERFACE but not validated by the KBA do not necessarily represent false positives. For example, INTERFACE implicates *DPEP1*, a peptidase, as a PCG for prolylglycine (C100003674), a dipeptide. As *DPEP1* is known to play a role in dipeptide hydrolysis [25], this pair is biologically plausible and demonstrates that INTERFACE can complement the KBA.

**Figure 5:**
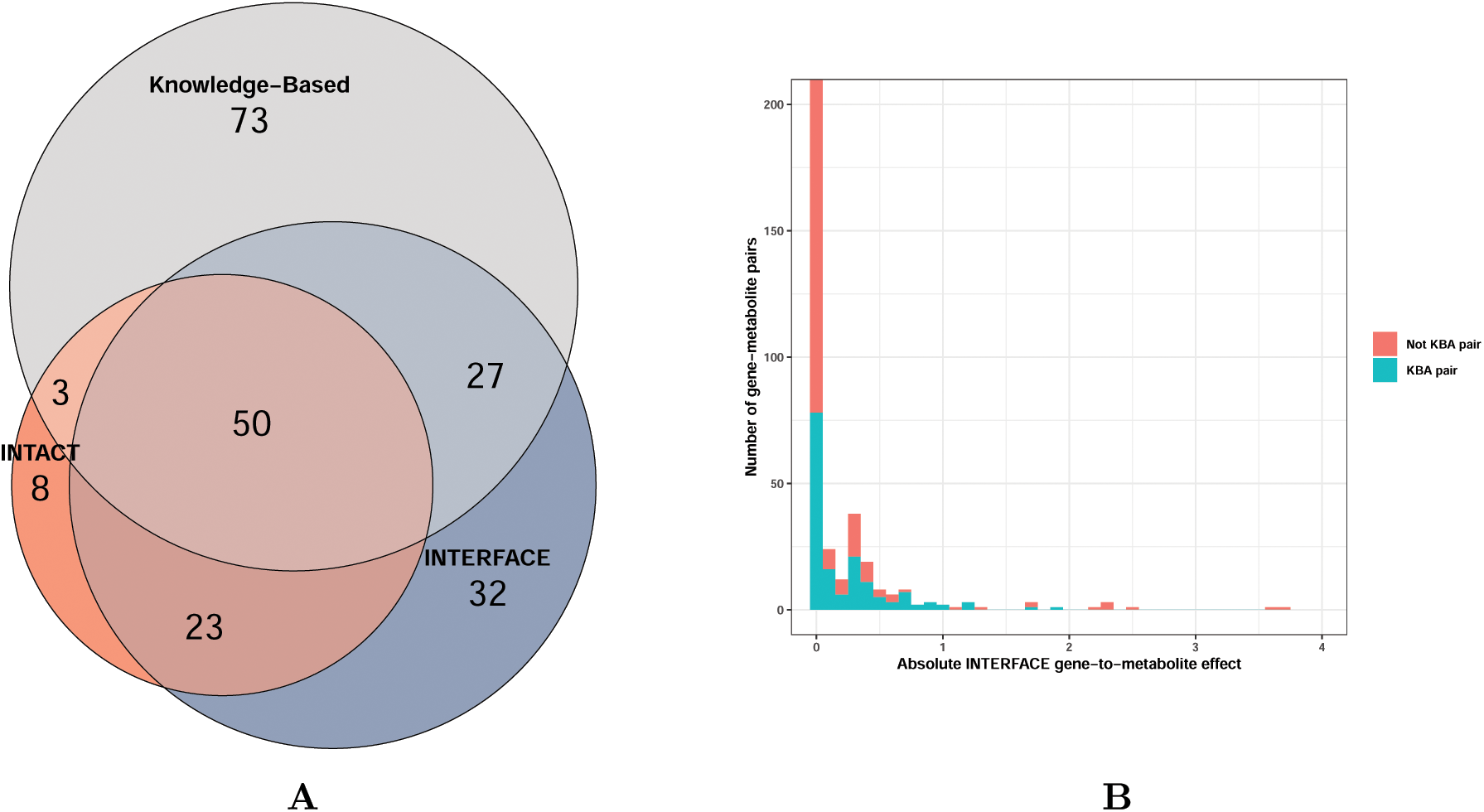
Comparison of PCG implication results by INTERFACE, INTACT, and knowledge-based approach from analyzing METSIM metabolite data. The left panel displays the overlap of gene-metabolite pairs implicated by the knowledge-based approach, INTERFACE, and INTACT, with the FDR control level at 0.05 applied to the INTERFACE and INTACT analyses. The right panel presents the histograms of the absolute INTERFACE gene-to-metabolite effect size estimates across all 5,509 tested gene-metabolite pairs. The distribution is stratified by whether the pair is validated by the KBA (KBA pair, in green) or not (Non-KBA pair, in red). The histogram is zoomed-in on the *y*-axis.

Additionally, we examine the gene-to-metabolite effects among pairs validated by the KBA. The right panel of the Figure 5 shows the distribution of absolute gene-to-metabolite effects estimated by INTERFACE for all tested gene-metabolite pairs. The mean absolute effect among all KBA-validated pairs is 0.230, which is significantly larger than the mean (0.007) for non-validated pairs (two-sample *t*-test *p*-value = 5.23 × 10^−14^). This analysis suggests that KBA results are enriched with causal genes with relatively large effects, and computational methods like INTERFACE can complement them by finding PCGs with modest effects. This analysis suggests that KBA results are enriched with causal genes that have relatively large effects, while computational methods like INTERFACE can complement these results by identifying PCGs with more modest effects.

To examine the specificity between the single-gene and multi-gene PCG implication methods, we focus on the set of 11 gene-metabolite pairs that are uniquely identified by INTACT but not INTERFACE. Notably, all 11 gene-metabolite pairs show moderate to strong colocalization evidence (gene-level colocalization probability, or GRCP, ≥ 0.50) and significant PWAS associations. However, upon closer inspection, all 11 cases are potentially false positives that fall into two categories.

The first category represents cases with highly inconsistent or heterogeneous gene-to-trait effect estimates across multiple independent instruments. An example of this kind is the gene *EPHX2*, located on chromosome 8 and identified by INTACT as a PCG for the unnamed metabolite C999918913. The gene shows both strong colocalization and PWAS evidence (GRCP = 1, PWAS *p*-value = 3.01 × 10^−6^) and a near certain INTACT posterior probability (= 0.989). However, the gene’s INTERFACE PIP is modest (0.468) and below the PCG discovery cutoff estimated at the 5% FDR control level.

We find that a single variant (rs2234887) is responsible for the strong colocalization signal. For this variant, PIPs in both the pQTL and GWAS analyses are nearly 1. However, the colocalized variant is only one of three independent pQTLs of *EPHX2* identified in the pQTL fine-mapping analysis by SuSiE, and the other pQTLs, also strong instruments, exhibit qualitatively different gene-to-trait effects (Figure 6). Specifically, rs2234887 suggests that increased *EPHX2* protein abundance decreases the target metabolite level, whereas the other two independent pQTLs indicate that increased protein abundance increases the metabolite level. The level of heterogeneity in these observed gene-to-trait effects, quantified by an *I*^2^ statistic [26], is 0.955 and close to the theoretical upper bound of 1. This evidence implies a potential violation of the IV/MR causality assumptions and reinforces the evaluation by INTERFACE. In total, we find that similar sensitivity analyses can explain 7 of the 11 gene-metabolite pairs in the INTACT-INTERFACE difference set. Details of these cases are provided in the Supplemental Figures S3-S8 and Supplemental Results.

**Figure 6:**
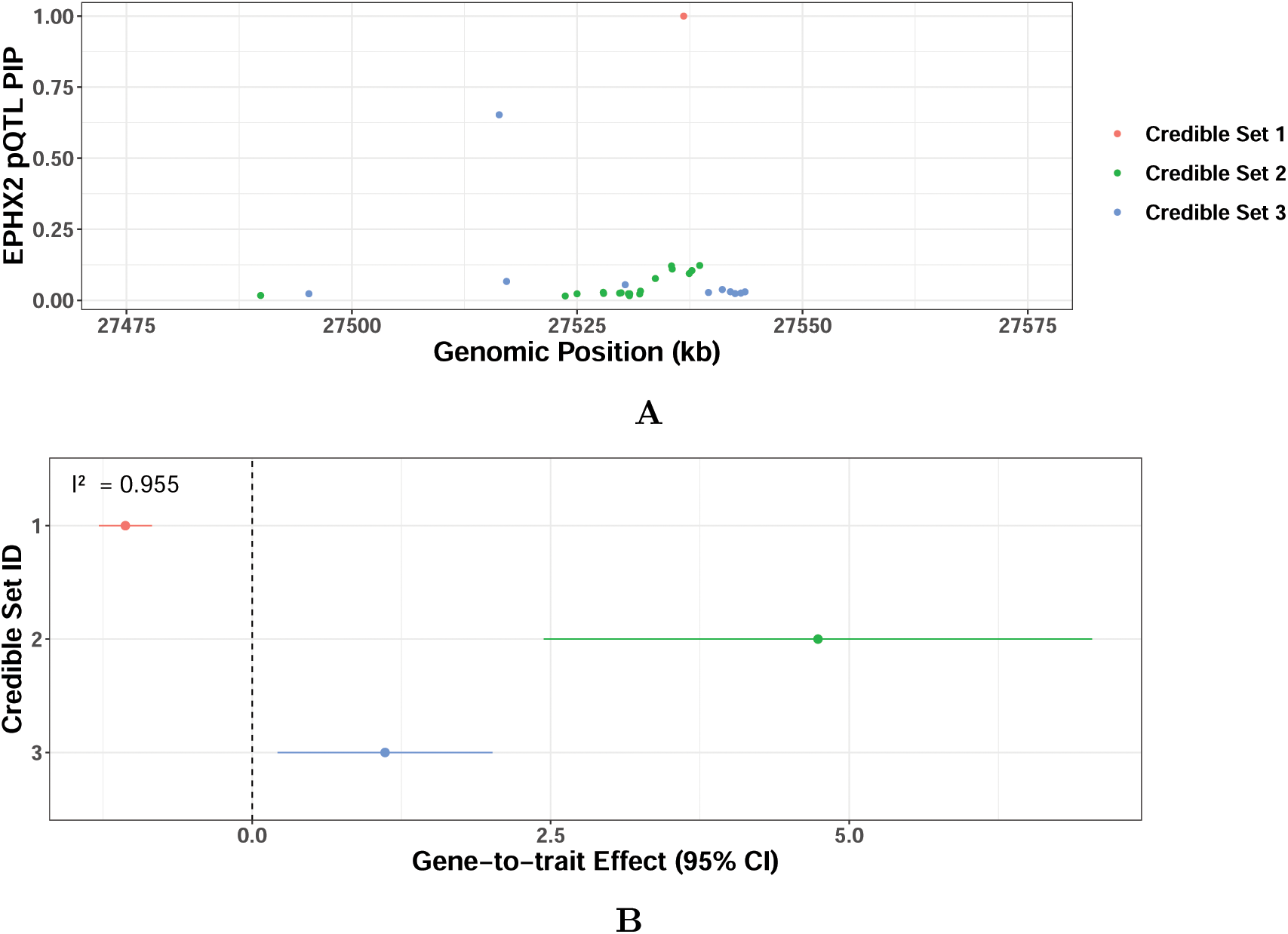
Sensitivity analysis for gene candidate. *EPHX2* The top panel (A) shows 3 independent pQTLs for *EPHX2* based on the SuSiE fine-mapping analysis of UK Biobank pQTL data. The SuSiE fine-mapping analysis of METSIM metabolite data in the region reveals only a single GWAS signal represented by a single SNP, rs2234887, completely overlapping with the first pQTL credible set. The bottom panel (B) presents the gene-to-trait effect estimates and the corresponding 95% confidence intervals by the top SNPs in the three pQTL credible set: rs2234887 (credible set 1), rs4149252 (credible set 2), rs751141 (credible set 3). While the colocalized SNP, rs2234887, suggests that increased protein levels lower the blood metabolite level of C999918913, the other two pQTLs suggest the opposite effect. The high degree of heterogeneity (*I*^2^ = 0.955) suggests potential violations of causal assumptions, which explains why INTERFACE does not assign a high PCG probability to *EPHX2*.

The second category concerns genes with correlated protein abundance. To illustrate, we focus on the gene *KLK15*, located on chromosome 19, which INTACT identifies as a PCG for the unnamed metabolite C999926054. This gene-metabolite pair has moderate colocalization and strong PWAS evidence (GRCP = 0.542, PWAS *p*-value = 4.87×10^−4^), but very low INTERFACE evidence (INTERFACE PIP *<* 0.001). In the same genomic region, another candidate gene, *KLK1*, shows similar colocalization evidence (GRCP = 0.524), but stronger PWAS association and INTERFACE evidence (PWAS *p*-value *<* 2.2 × 10^−16^, INTERFACE PIP = 0.998). Upon close inspection, we find that the two genes share a pQTL responsible for the colocalization signal, and their genetically predicted protein abundance levels are modestly correlated (*ρ* = 0.43, p-value *<* 2.2 × 10^−16^). The complete pattern of correlations for all genes in this region is shown in Supplemental Figure S9. As a result, both genes are implicated by the single-gene analysis method, assessed with similar levels of evidence. However, when they are simultaneously analyzed by INTERFACE, the association evidence of *KLK15* appears to be explained away by *KLK1*. For confirmation, we fit a multi-gene PWAS model by removing the region genotype matrix, which accounts for direct GWAS effects, from the INTERFACE model. This analysis yields results similar to the INTERFACE finding (PIPs for *KLK15* and *KLK1* are *<* 0.001 and 1.00, respectively). Additionally, when we remove the predicted protein level of *KLK1* from the original INTERFACE model, we observe a substantial increase in the *KLK15* PIP to 0.226. Although our analysis does not rule out the biological relevance of *KLK15* for the target metabolite, INTERFACE recognizes that there is only a single causal gene within the region and *KLK1* is more likely to be the sole PCG based on observed data.

## 3 Discussion

This paper introduces INTERFACE, a computational method for fine-mapping putative causal genes of complex traits by integrating functional molecular QTL data. INTERFACE assumes a structural equation model compatible with commonly applied instrumental analysis, Mendelian randomization, and TWAS methods. By considering multiple genes within a genomic region and explicitly controlling direct variant-to-gene effects, the proposed SEM covers a more comprehensive set of scenarios than most existing causal gene implication methods. Through extensive simulations, we demonstrate that INTERFACE improves specificity and sensitivity for detecting PCGs and provides accurate gene-to-trait effect estimates. When used to analyze METSIM metabolite and UK Biobank pQTL data, INTERFACE fine-maps 57% more PCGs than the existing single-gene PCG implication method. Additionally, a substantially higher number of PCGs implicated by INTERFACE are independently validated by a knowledge-based approach. We also observe that INTERFACE’s findings complement and expand existing knowledge of metabolic pathways, highlighting the benefits of data-driven discoveries.

Due to computational constraints, running INTERFACE at the genome or chromosome scale is currently impractical. Therefore, we recommend using single-gene PCG implication methods to screen and identify genomic regions for subsequent INTERFACE analysis. The overall procedure, as demonstrated in our analysis of METSIM metabolite data, is analogous to the common practice of genome-wide genetic association studies, where single-SNP analysis serves as a preliminary screening step, followed by multi-SNP fine-mapping analysis in selected regions.

The prior specification is a crucial component of the INTERFACE model. In the Methods section, we demonstrate that INTERFACE can include several existing methods as special cases by adjusting the priors for the SEM (1), leading to significantly different performance outcomes. The INTERFACE prior is uniquely advantageous because it is interpretable, justifiable, and data-informed. Specifically, the formulation of the PCG prior, which combines colocalization and TWAS evidence, can be understood as an application of Dempster-Shafer decision theory [11, 16, 27]. In simulations, we observe that the INTERFACE with the proposed prior specification achieves near-optimal performance.

Probabilistic quantification of statistical evidence has become increasingly common with applications of Bayesian methods in genetic association analysis. INTERFACE leverages probabilistic evidence from genetic association and integrative analyses as input and evaluates the plausibility of a candidate gene being a PCG by assessing a posterior probability. These PCG probabilities enable rigorous Bayesian or local false discovery rate control procedures, critical for evaluating and controlling false positive findings. In contrast, a frequentist approach may struggle in this context, as PCG fine-mapping is inherently a variable/model selection problem rather than a hypothesis testing problem.

We acknowledge that our proposed computational method and data analysis do not come without limitations. First and foremost, integrative analysis is fundamentally constrained by the power of genetic association analyses for molecular and complex traits. Despite the recent efforts to increase sample sizes to the biobank scale, our ability to uncover genetic associations remains modest. Our simulations intentionally generate data matching observed heritability from METSIM metabolite and UK Biobank pQTL studies. Even with state-of-the-art statistical fine-mapping approaches, we find that a significant proportion of true genetic association signals are not identified. Since GWAS results serve as imperfect inputs for subsequent integrative analyses, the overall power for discovering PCGs is understandably affected. A similar phenomenon has also been discussed in the context of colocalization analysis [14]. Second, like all causal inference methods based on observational data, the causal interpretation of INTERFACE results relies on a set of assumptions similar to those in instrumental variable/Mendelian randomization analysis. Although INTERFACE has built-in functionality for automatic sensitivity diagnosis to detect severe violations of causal assumptions (Section 2.2), its success largely depends on the observed data, such as the availability of independent molecular QTL instruments and the level of heterogeneity in their corresponding gene-to-trait effect estimates. Therefore, we recommend that practitioners carefully interpret INTERFACE results and seek corroborating evidence from independent knowledge sources and experiments. Third, our analysis of METSIM metabolites exclusively uses UK Biobank *cis*-pQTLs due to concerns about the quality and reliability of available *trans*-pQTLs obtained through current proteomic technologies [28, 29]. However, the proposed model and fitting algorithm are fully capable of utilizing high-quality *trans* molecular QTLs, which can serve as reliable, independent instruments to improve both PCG discovery and gene-to-trait effect estimation. Fourth, the current version of INTERFACE does not infer the molecular QTLs’ relevance to tissue or cell type. However, the output from tissue-specific applications of INTERFACE, particularly the gene-to-trait effect estimates, may provide valuable information and a starting point to address this longstanding issue.

Finally, future work can further improve the INTERFACE method. First, the INTERFACE model could be expanded to simultaneously consider multiple molecular phenotypes directly linked to gene functions, such as expression levels, isoform usage, and protein abundance. Recent work [30] has demonstrated that incorporating information from multiple gene products can significantly improve the power of PCG discovery in single-gene analyses. Second, a wealth of genomic and epigenomic data, such as various types of gene networks and gene regulatory annotations, could be leveraged for PCG discovery. These additional information sources can naturally be incorporated as priors within the INTERFACE framework.

## 4 Methods

### 4.1 Model details

The SEM model (1) accounts for unobserved confounding shared in ***e****_M,i_* and ***e****_Y_* when applied in a one-sample design, where all molecular and complex-trait phenotypes are measured from the same samples. Consequently, ***M*** *_i_* is correlated with ***e****_Y_* and becomes an endogenous variable. To mitigate this correlation for proper inference of *γ_i_*, a common approach is to replace ***M*** *_i_* with its predicted value from genetic instruments,i.e., ***M̂*** *_i_*. The inference is then based on the working model (2). Notably, Model (2) is also applicable in two-sample designs, where the samples used to measure molecular phenotypes and construct ***M̂*** *_i_* do not overlap with those used to measure complex-trait phenotypes.

Our primary objective is to infer the *γ_i_* values. To determine if the target gene *i* is a PCG, we aim to compute the posterior inclusion probability, Pr(*γ_i_* ≠ 0 | Data). Additionally, we seek to estimate the magnitude of the corresponding gene-to-trait effect, which can then be used to rank PCGs.

### 4.2 Prior specifications

The three prior distributions critical for INTERFACE inference are

- The prior probability that the region of interest has neither a PCG nor a direct-effect SNP, i.e., Pr(***γ*** = **0** and ***β****_Y_* = **0**)
- The prior probability that SNP *k* has a direct effect on the complex trait, i.e., Pr(*β_Y,k_* ≠ 0)
- The prior probability that gene *i* is a PCG, i.e., Pr(*γ_i_* ≠ 0)

These priors are informed by the results from fine-mapping, colocalization, and single-gene TWAS analyses with respect to the complex and molecular traits. Specifically, these analyses provide key variant- and gene-level estimates, including:

1. The proportion of genome-wide variants that are GWAS hits, *p̂_g_*
2. The proportion of genome-wide variants that are QTLs, *p̂_m_*
3. The proportion of genome-wide variants that are colocalized, *p̂_c_*
4. The proportion of genome-wide genes that are TWAS genes, *π̂*
5. The gene-level colocalization probability for each candidate gene *i*, GRCP*_i_*

These estimates are derived from existing computational methods [5, 16, 31, 32] and widely used in various genetic association and integrative analyses. A summary of the corresponding estimation methods is provided in the Supplemental Methods.

Based on SEM (1), a necessary and sufficient condition for a genomic region to contain neither a PCG nor a direct-effect SNP is the absence of GWAS hits in that region. Hence, we set

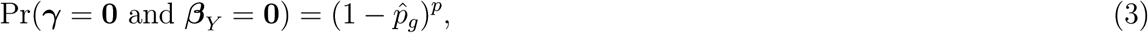

where *p* denotes the number of SNPs in the genomic region of interest.

The prior probability that a variant has a direct effect can be derived from the proportion of SNPs that are only GWAS hits but not molecular QTLs, i.e.,

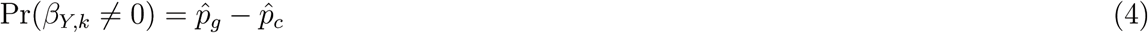

For the PCG prior, we follow the logic of INTACT [16] and use a multiplicative combination of colocalization evidence and TWAS prior information, i.e.,

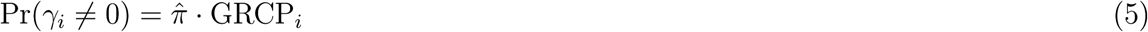

The following rationale justifies this formulation. First, colocalization is a necessary condition for a causal gene, as mathematically implied by (1). However, colocalization does not establish causality due to the potential for pleiotropy. Thus, the prior probability that a gene is causal should be bound by its colocalization evidence, i.e.,

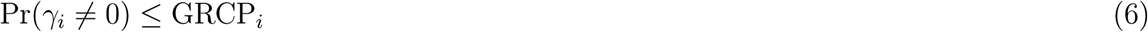

Second, in an ideal scenario where colocalization evidence for each gene is near perfect (i.e., GRCP*_i_* → 0 or GRCP*_i_* → 1, for all genes), *π̂*, estimated from the TWAS *p*-value distribution using Storey’s method, serves as a natural conservative prior for each candidate gene. Third, the multiplicative combination of GRCP*_i_* and *π̂* can be justified from the perspective of the evidence combination rule in Dempster-Shafer theory [27], where we utilize *a priori* probabilistic evidence from colocalization and TWAS to assess the putative causality of a gene candidate. As demonstrated by Hukku et al. [11], the genes implicated by colocalization and TWAS analyses are largely non-overlapping in practice, suggesting that their reliability in inferring PCGs is approximately DS-independent, thereby justifying the use of Dempster’s rule of combination in (5).

The prior specification is an integral part of the INTERFACE model. It distinguishes it from the existing methods, many of which can be seen as special cases of the INTERFACE model with alternative prior specifications. For example, setting Pr(*β_Y,k_* = 0) = 1 for all SNPs removes the possibility of direct variant-to-trait effects from consideration. This approach makes the resulting method susceptible to false positives by failing to control for LD hitchhiking/indirect horizontal pleiotropy explicitly. Most existing single-gene and multi-gene TWAS methods [1, 2, 3, 33] fall into this category. Among the methods that explicitly consider direct-effect SNPs, FOCUS [10] assumes all SNPs in a region have the same constant direct effect on the trait of interest, similar to the burden test assumption commonly used for detecting rare-variant associations. The prior form for this assumption is discussed in [34]. However, the assumption seems too strong for common variant associations. In practice and our simulation studies, this approach has been shown to have limited ability to control LD hitchhiking [16]. Another näıve prior specification is to assign equal/exchangeable priors for Pr(*γ_i_* ≠ 0) and Pr(*β_Y,k_* ≠ 0) for all genes and SNPs. The resulting algorithm is equivalent to running SuSiE directly to fit the model (2) under its default setting. Since the frequencies of causal genes and SNPs can differ by orders of magnitude in many complex traits, this prior specification can lead to highly conservative results for implicating PCGs. The recently proposed cTWAS method also uses an empirical Bayes procedure to estimate two key prior quantities, Pr(*γ_i_* ≠ 0) = *P̂*_gene_ and Pr(*β_Y,k_* ≠ 0) = *P̂*_snp_, from the observed data. However, to overcome computational challenges, their EM procedure requires additional data pruning and introduces further simplifying assumptions, such as limiting each gene to at most one *cis* molecular QTL. Additionally, they set Pr(***γ*** = **0** and ***β****_Y_* = **0**) = 1 − *P̂*_snp_− *P̂*_gene_, which appears overly conservative in most practical scenarios. The common drawback of these alternative prior specifications is that they all implicitly or explicitly make rather strong assumptions without support from observed data or existing knowledge.

### 4.3 Estimating gene-to-trait effects

We examine the analytical factors that impact the accuracy and reliability of gene-to-trait effect estimates in INTERFACE.

#### 4.3.1 Instrument selection using QTL fine-mapping results

The importance of selecting strong instruments for causal effect estimation is well-established and extensively researched in the literature. The signal-to-noise ratios of the corresponding instrument-exposure association evidence are typically used to quantify the strength of the instruments. For example, the *F* -statistic of a candidate instrument is commonly used to measure the instrument strength in a regression model. In our context, the widespread linkage disequilibrium (LD) complicates the selection of a single best variant to represent the underlying causal molecular QTL. State-of-the-art fine-mapping methods [18, 31] report a credible set or signal cluster comprising multiple likely causal variants, along with their posterior inclusion probabilities (PIPs), for each identified molecular QTL. We propose using the cumulative PIP (CPIP) of a signal cluster or the coverage probability of a credible set as a metric to quantify the strength of the corresponding genetic instruments. The CPIP value reflects the signal-to-noise ratio of an underlying genetic association signal and has the unique advantage of accounting for variant-level uncertainty due to LD, making it better suited for measuring the overall strength of a molecular QTL. For a given CPIP threshold, we only consider member SNPs of a signal cluster or credible set that exceeds the threshold to predict the corresponding molecular phenotype.

#### 4.3.2 Constructing TWAS prediction models

We make a straightforward modification to the PTWAS method [3] to incorporate the above instrument selection criterion when constructing TWAS prediction models. Specifically, for a given PTWAS prediction model, we shrink the weights of genetic variants excluded from the qualified signal clusters or credible sets to exactly zero. This simple procedure ensures that only the selected instruments contribute to the molecular phenotype prediction.

For molecular fine-mapping results generated by SuSiE, as in our UK Biobank pQTL data analysis, we treat the reported posterior means of all variants as the raw TWAS weights and apply a similar shrinkage procedure. In our simulations, we observe that this approach produces prediction models that are numerically nearly identical to the original PTWAS construction.

In our numerical experiments and data analysis, we also apply two other popular TWAS methods: PrediXcan and SMR.

PrediXcan [2] uses elastic net regularization [35] to construct molecular trait prediction models. This method regularizes potential QTL effects from all SNPs, effectively selecting only the strongest instruments. The degree of shrinkage by the elastic net is determined through internal cross-validation based on prediction performance. Our simulations show that PrediXcan achieves the highest power for detecting PCGs among all the prediction models considered. However, its gene-to-trait effect estimates exhibit some systematic bias, likely due to the universal shrinkage applied to TWAS weights. Despite this, we do not observe severe outlying estimates of gene-to-trait effects with PrediXcan models.

We also apply a version of SMR that estimates the gene-to-trait effect using the statistically most significant QTL SNP. SMR is equivalent to building a prediction model with only a single SNP [4, 11]. While it performs adequately with underpowered molecular QTL data, it is often outperformed by methods that can effectively utilize multiple independent instruments [2, 3].

SMR can also be viewed as a strong form of screening or regularization on genetic instruments, where all weights are shrunk to zero except for the most significant QTL SNP. In our simulations, SMR does not produce outlying gene-to-trait effect estimates, but it tends to underestimate gene effects, which is reflected in its reduced power to detect PCGs.

### 4.4 Model-fitting with variational inference algorithm

We apply the Sum of Single Effects (SuSiE) algorithm [18] to fit the model (2) to the observed data. SuSiE is an efficient variational inference algorithm designed for solving the variable selection problem in linear regression models. In addition to specifying INTERFACE-specific priors, we set the *L* parameter, which represents an upper bound for the total number of direct-effect SNPs and PCGs, to a default value of 10. For output, we focus on the estimated PIPs for all candidate genes, as well as the posterior mean and variance for the corresponding gene-to-trait effects. We also experiment with an alternative algorithm, DAP-G, to fit the model (2) and observe nearly identical numerical results for PIP estimation.

### 4.5 Simulation data generation and processing

We design simulations to evaluate INTERFACE’s performance, focusing on its ability to identify PCGs in the presence of direct-effect SNPs. We select a 3 Mb genomic region on chromosome 3, containing 39 protein-coding genes and 26,986 SNP genotypes from 706 GTEx samples. This region has been highlighted by Hukku *et al.* [11], who demonstrated that a single direct-effect variant, rs2871960, leads to extensive spurious TWAS signals within the region when jointly analyzing UK Biobank height and GWAS Whole Blood eQTL data. The LD patterns in this region are complex but not uncommon (Supplementary Figure 1).

In all simulated datasets, we designate SNP rs2871960 as the sole true direct-effect SNP. For each of the 39 genes, we randomly select two distinct *cis*-eQTLs. We randomly draw either 0, 1, or 2 causal genes in each simulation with probabilities of 0.5, 0.25, and 0.25, respectively. As a result, approximately half of the simulated datasets represent the null scenario. Given the eQTLs, causal genes, and the direct-effect SNP, we then generate complex trait data and each gene’s molecular phenotype data using the SEM (1). The genetic effects for all the eQTLs, the direct variant-to-trait effects, and the gene-to-trait effects are independently drawn from a N(0, 0.6^2^) distribution. The residual error for each trait and each sample is independently drawn from a standard normal distribution. We simulate 600 datasets based on this scheme. The mean (SD) proportion of variance explained (PVE) for molecular trait levels across all genes is 0.16 (0.13), while the mean (SD) PVE for complex trait levels across all datasets is 0.14 (0.13). It is important to note that the data-generative scheme in our simulations is not compatible with the INTERFACE model: The generative model for the simulation assumes exchangeable priors among causal genes, which is more akin to the cTWAS model.

For pre-processing, we perform separate probabilistic fine-mapping analyses on each simulated dataset for both complex and molecular traits using DAP-G and SuSiE. These fine-mapping results are subsequently used for colocalization and TWAS analyses. Specifically, we treat the 600 simulated regional datasets as 600 fine-mapping target regions from a single integrative analysis. This approach allows us to pool all simulated data together to estimate the global parameters, *p̂_g_*, *p̂_m_*, and *p̂_c_*, as the by-products of colocalization enrichment analysis using fastENLOC (and similarly estimate *π̂* from the TWAS results). We then compute the necessary priors for each simulated dataset and perform the INTERFACE analysis.

For the simulated data, the estimated values of *p̂_g_*, *p̂_m_*, and *p̂_c_* are 3.37 × 10^−5^ (3.71 × 10^−5^), 1.58 × 10^−3^ (2.83 × 10^−3^), and 1.25 × 10^−5^ (5.56 × 10^−5^), respectively. The numbers in parentheses represent the true frequencies. It is known that the multiple imputation scheme implemented in fastENLOC, combined with the imperfect power of genetic association analysis, tends to produce conservative estimates for these quantities [36, 14]. However, the estimates are generally close to the true frequencies and conservative. The estimated *π̂* from the PTWAS analysis is 0.274, which is significantly higher than the true proportion of causal genes (0.019). Nevertheless, after incorporating colocalization evidence, the estimated INTERFACE gene priors become much more conservative with the mean value = 0.007 (Supplemental Figure S2).

### 4.6 Methods in comparison

Using the simulated data, we compare INTERFACE with two classes of integrative analysis methods.

The first class consists of single-gene analysis methods, including PrediXcan [2], SMR [4, 11], PTWAS [3], fastENLOC [5, 14], and INTACT [16]. PrediXcan, SMR, and PTWAS are TWAS methods that apply different analytical strategies to construct genotype-based molecular phenotype prediction models. Consequently, they exhibit varying levels of power in detecting PCGs and differ in the accuracy of their gene-to-trait effect estimates. However, all these methods are susceptible to false positives due to LD hitchhiking [11]. fastENLOC is a colocalization method that leverages probabilistic fine-mapping analysis of both molecular and complex traits but suffers from a fundamental lack of power [14]. INTACT integrates both TWAS and colocalization evidence, achieving a balance between the sensitivity of TWAS methods and the specificity of colocalization approaches.

The second class analyzes multiple genes within a genomic region simultaneously. From the perspective of INTERFACE, each method in this class can be derived from (2) with an alternative prior specification. Multi-gene TWAS assigns an equal prior of 1*/q* to each of the *q* genes, assuming no direct variant-to-trait effects (i.e., *P* (***β****_Y_* = 0) = 1) in the genomic region of interest. The “SuSiE default” method assumes an exchangeable prior of 1*/*(*p* + *q*) across all genes and SNPs, meaning that it does not distinguish between gene-to-trait effects (***γ***) and the direct variant-to-trait effects (***β****_Y_*) *a priori*. The FOCUS method [10] also assigns an equal prior of 1*/q* to all genes, while its treatment of potential direct variant-to-trait effects is equivalent to using the following structured flat prior,

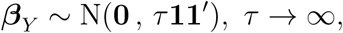

which enforces equal effect size for all SNPs [34]. The cTWAS method estimates Pr(*γ_i_* ≠ 0) = *P̂*_gene_ and Pr(*β_Y,k_* ≠ 0) = *P̂*_snp_ under certain simplifying assumptions, and specifies Pr(***γ*** = **0** and ***β****_Y_* = **0**) = 1−*P̂*_snp_−*P̂*_gene_. Finally, the oracle prior uses the true exchangeable probabilities for ***γ*** and ***β****_Y_* from our generative model for the simulation.

### 4.7 Analysis of METSIM Metabolon metabolite GWAS and UK Biobank pQTL data

The METSIM Metabolon metabolite study involves 10,188 Finnish men from Kuopio, examined between 2005 and 2010 [37]. Whole-genome sequencing of the study participants yielded over 26 million variants that passed quality control procedures. We first identified index SNPs through single-variant and stepwise conditional analyses in the genetic association analysis. We then defined genomic regions for fine-mapping by extending 1 Mb on either side of the index SNPs. Subsequently, we conducted multi-SNP fine-mapping analyses for these regions using SuSiE [18], with association evidence summarized by SuSiE 95% credible sets. A comprehensive description of the METSIM metabolite data pre-processing and genetic association analysis is provided in the Supplemental Methods.

We obtained UK Biobank pQTL data as processed summary information, where each inferred *cis*-pQTL is represented by a SuSiE 95% credible set. The member SNPs, the corresponding PIPs, and the posterior mean effect are reported within each credible set. We pool the SuSiE pQTL credible sets across all examined proteins to generate the required probabilistic annotations for colocalization analysis using fastENLOC. For PWAS analysis, we follow the modified PTWAS procedure, constructing weights for the predicted protein levels of a target protein using the posterior mean effects of all member SNPs in the available credible sets.

After running colocalization and PWAS analyses, we proceed to select region-metabolite pairs for INTERFACE analysis. The selection procedure involves three steps:

1. Merge overlapping *cis*-fine-mapped regions for 2,040 UK Biobank proteins into 293 distinct regions.
2. Apply INTACT to perform a pQTL-GWAS integrative analysis, flagging PCGs at the 25% FDR level for all metabolites.
3. Select the region containing at least one flagged PCG for INTERFACE analysis.

We select 193 unique region-metabolite pairs for PCG fine-mapping using the above procedure.

In the INTERFACE analysis of each selected genomic region, we include the predicted protein levels of all genes and all common SNPs with a minor allele frequency greater than 3%. The INTERFACE priors are computed using the estimates of *p̂_g_*, *p̂_m_*, and *p̂_c_* from fastENLOC and *π̂* from PTWAS *p*-values.

