## Supplemental Figures and Methods for "Probabilistic Fine-mapping of Putative Causal Genes"

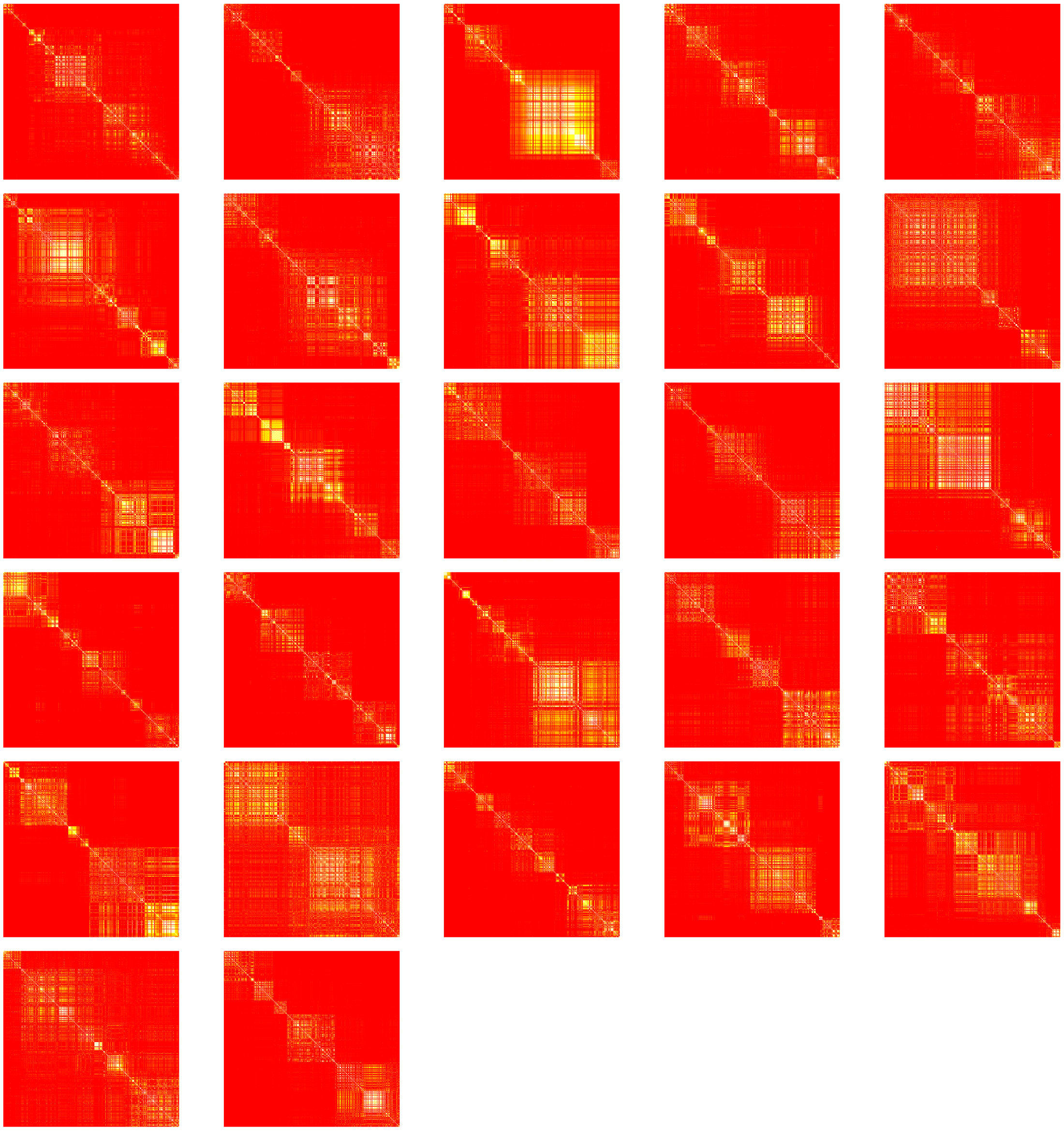

Figure S1: LD structure of the chromosome 3 region used in the simulation study. The 26,986-SNP region is divided into 27 blocks of approximately 1,000 SNPs each. LD is measured by pairwise $r^2$. Blocks are ordered by position, from left to right, top to bottom. The plots represent 706 individuals from the GTEx skeletal muscle data.

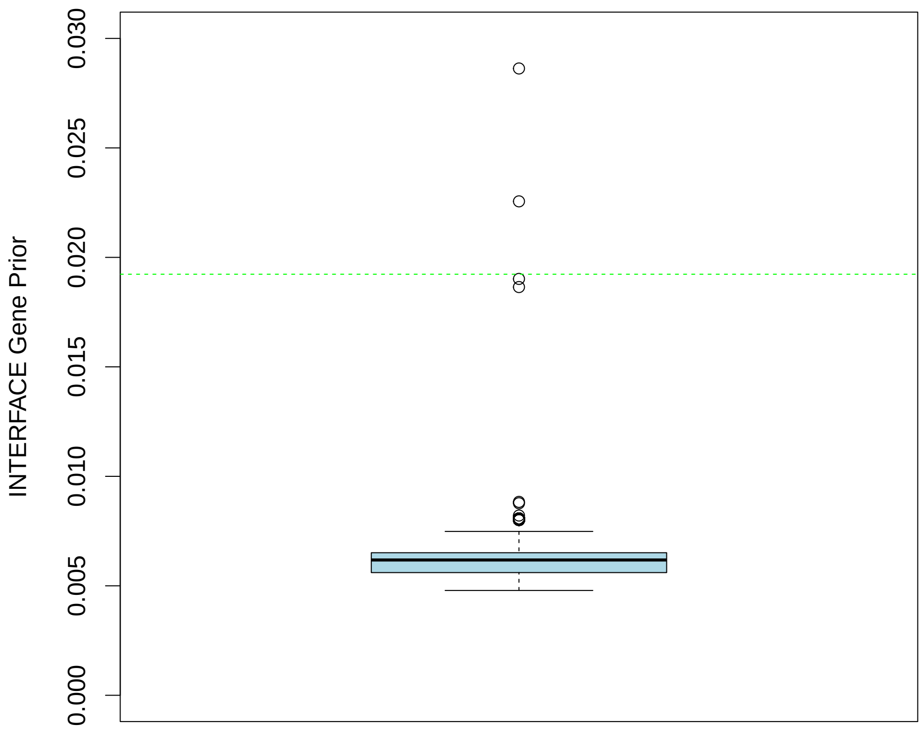


Figure S2: Boxplot of estimated INTERFACE gene priors in the simulated dataset. The green dotted line represents the true frequency of causal genes in the dataset (0.019). The plot is truncated at 0.03, and approximately 0.4% of the simulated genes have estimated priors greater than 0.03. The maximum estimated prior is 0.268, which is close to the estimated $\hat{\pi}$ value of 0.274.

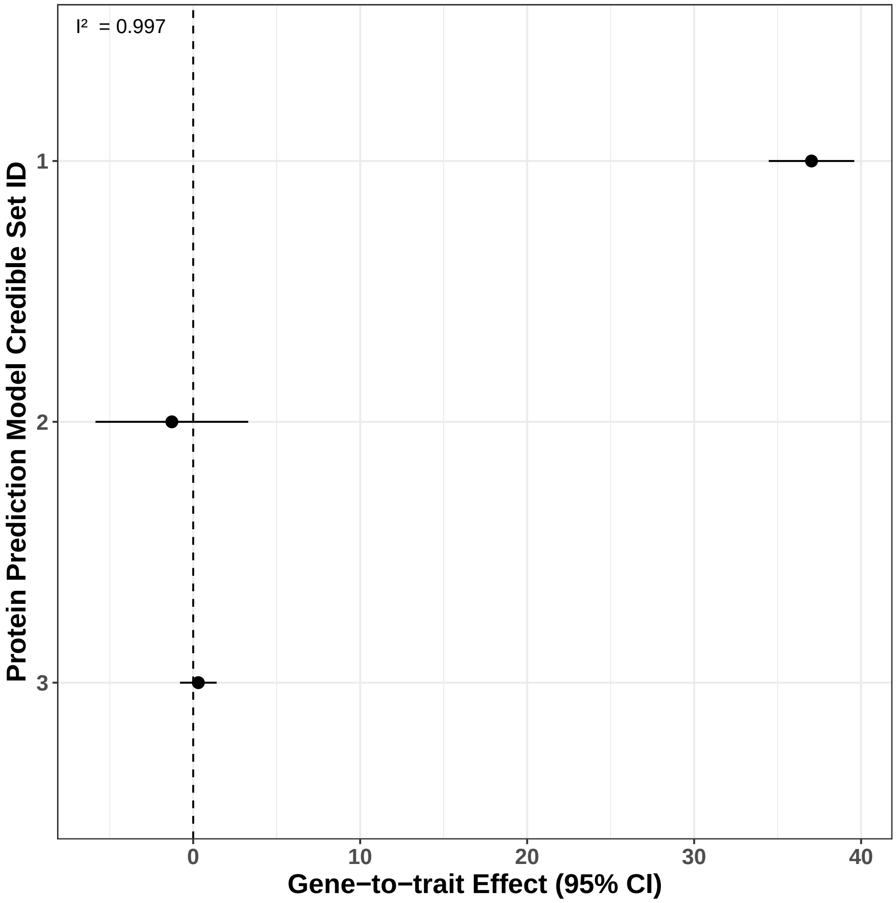


Figure S3: VGF-to-C100001293 INTERFACE effect estimates based on single-varant prediction models. The y-axis denotes the pQTL credible set used to predict VGF protein levels. For each of the three independent credible sets, we use the variant with the highest PIP in a single variant prediction model. We fit a simple linear regression model three times, changing the VGF predicted protein levels each time. The x-axis denotes the gene-to-trait effect estimate and 95% confidence interval. The $I^2$ statistic quantifies the heterogeneity of the independent effect estimates.

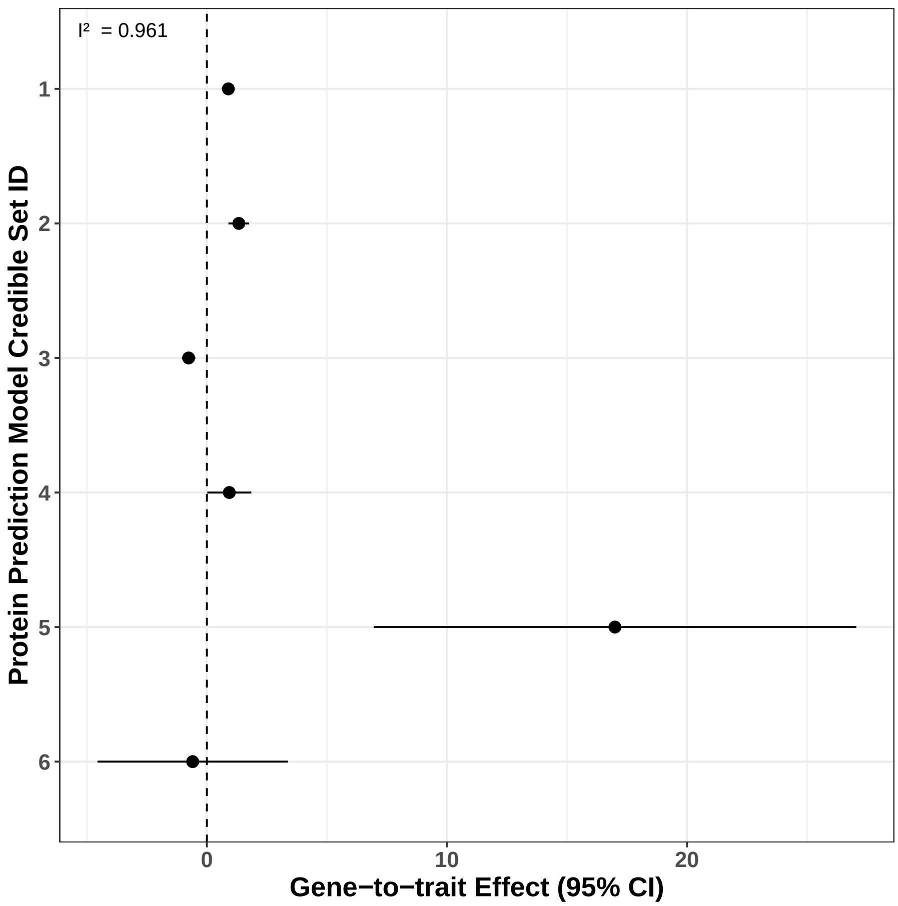


Figure S4: The AOC1-to-C100004541 INTERFACE effect estimates based on single-varant prediction models. The y-axis denotes the 6 pQTL credible set used to predict AOC1 protein levels. For each of the six independent credible sets, we use the variant with the highest PIP in a single variant prediction model. We fit a simple linear regression model six times, changing the AOC1 predicted protein levels each time. The x-axis denotes the gene-to-trait effect estimate and 95% confidence interval. The $I^2$ statistic quantifies the heterogeneity of the independent effect estimates.

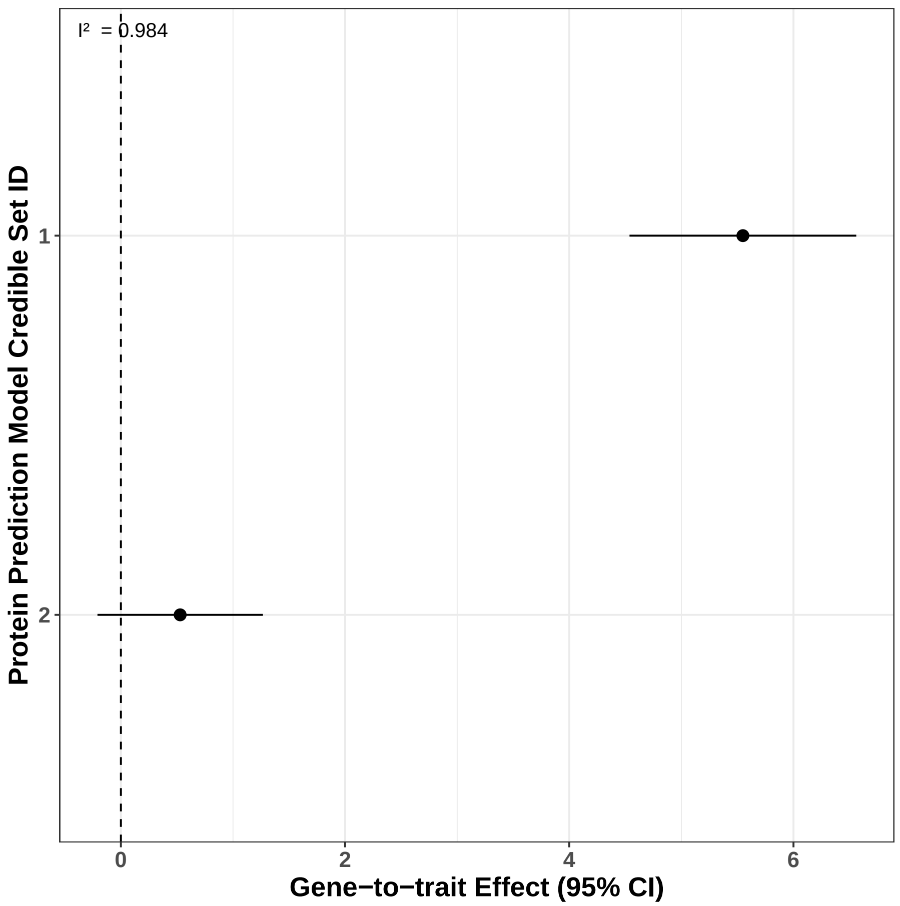


Figure S5: The CHMP1A-to-C100021198 INTERFACE effect estimates based on single-varant prediction models. The y-axis denotes the 2 pQTL credible set used to predict CHMP1A protein levels. For each of the two independent credible sets, we use the variant with the highest PIP in a single variant prediction model. We fit a simple linear regression model twice, changing the CHMP1A predicted protein levels each time. The x-axis denotes the gene-to-trait effect estimate and 95% confidence interval. The $I^2$ statistic quantifies the heterogeneity of the independent effect estimates.

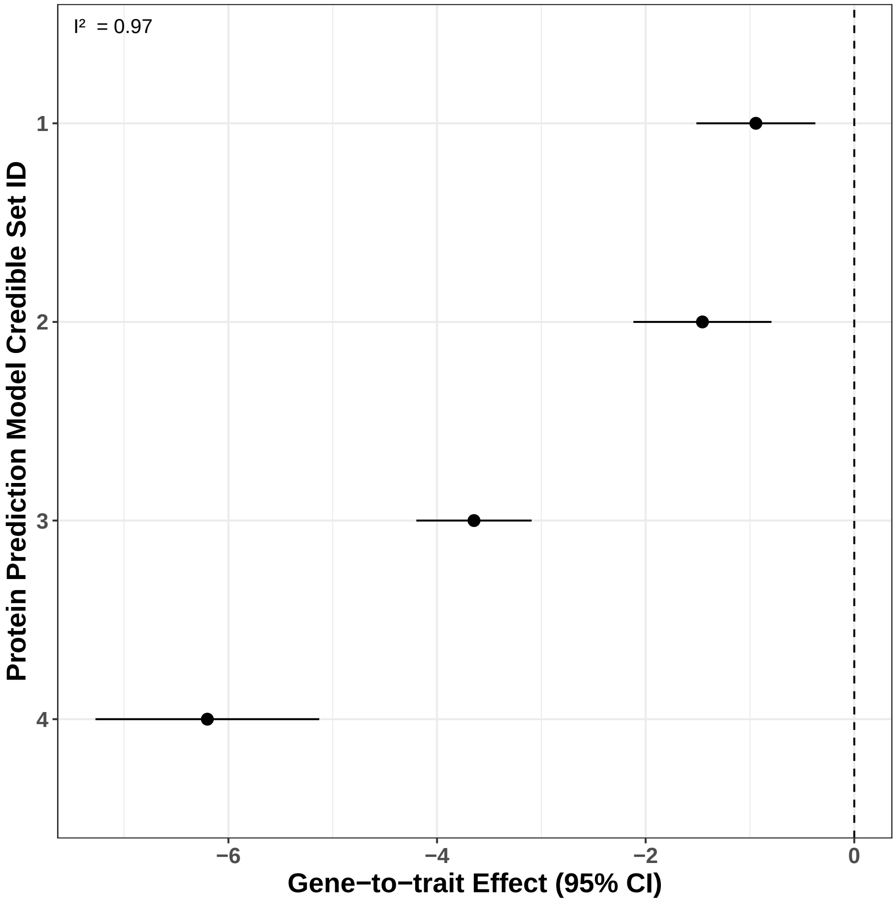


Figure S6: The OPLAH-to-C1021 INTERFACE effect estimates based on single-varant prediction models. The y-axis denotes the 4 pQTL credible set used to predict OPLAH protein levels. For each of the four independent credible sets, we use the variant with the highest PIP in a single variant prediction model. We fit a simple linear regression model four times, changing the OPLAH predicted protein levels each time. The x-axis denotes the gene-to-trait effect estimate and 95% confidence interval. The $I^2$ statistic quantifies the heterogeneity of the independent effect estimates.

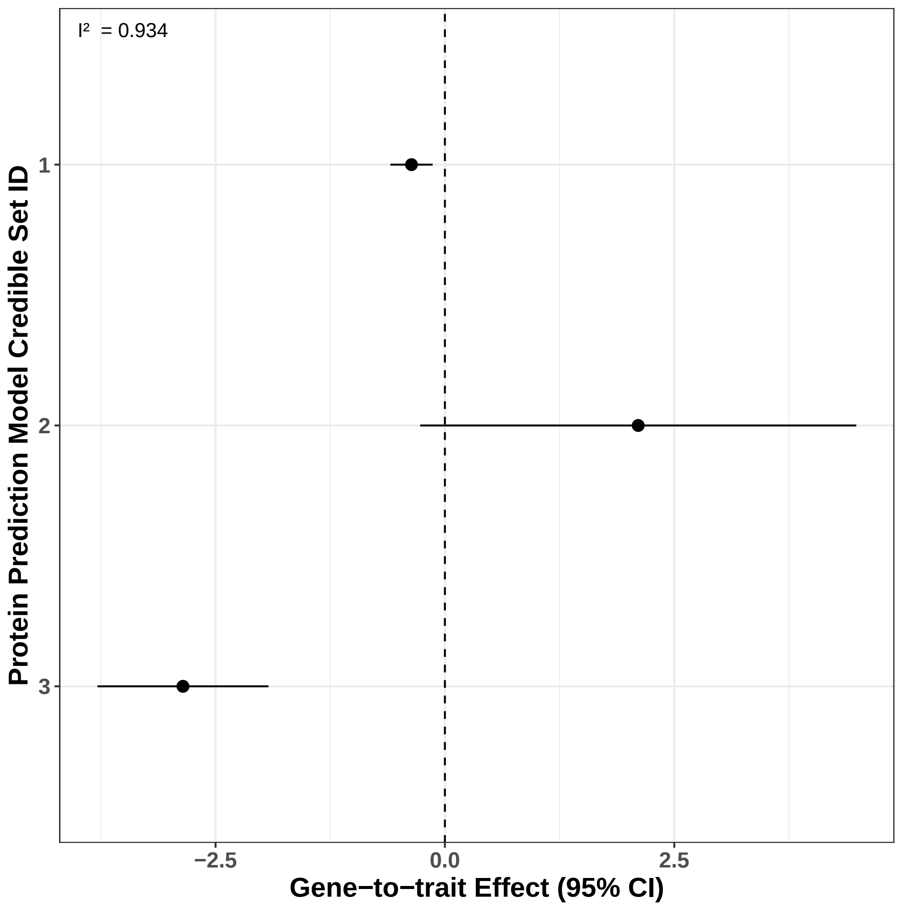


Figure S7: The EPHX2-to-C999918899 INTERFACE effect estimates based on single-varant prediction models. The y-axis denotes the 3 pQTL credible set used to predict EPHX2 protein levels. For each of the three independent credible sets, we use the variant with the highest PIP in a single variant prediction model. We fit a simple linear regression model three times, changing the EPHX2 predicted protein levels each time. The x-axis denotes the gene-to-trait effect estimate and 95% confidence interval. The $I^2$ statistic quantifies the heterogeneity of the independent effect estimates.

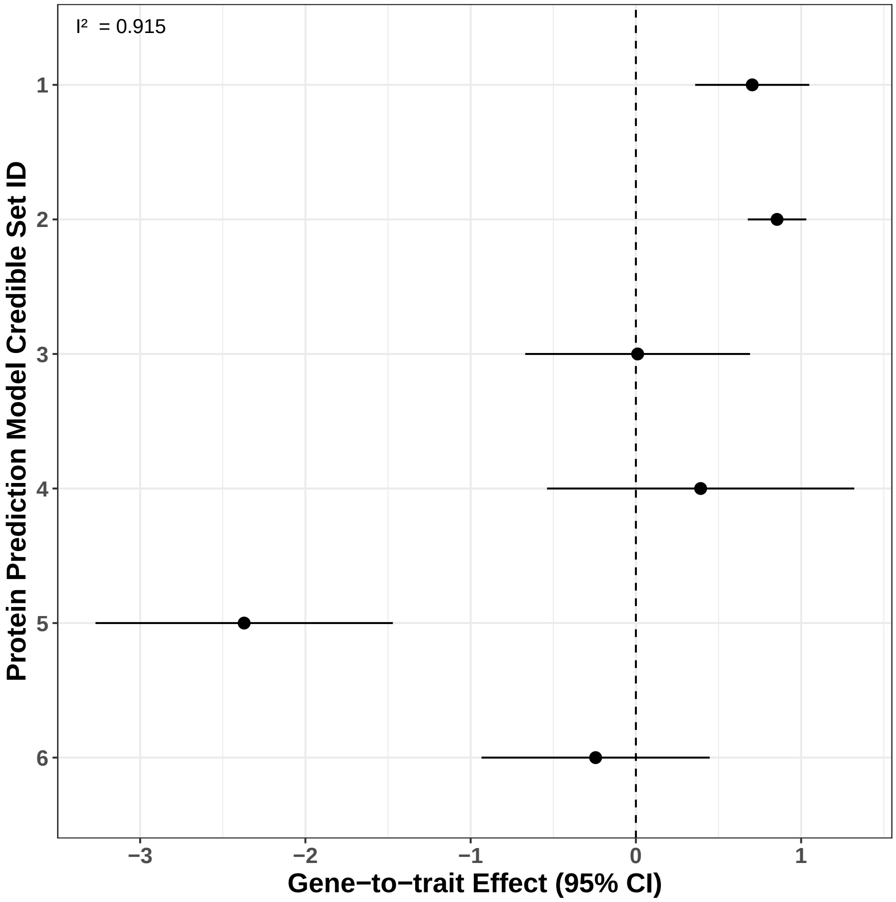


Figure S8: The TIMP4-to-C999924328 INTERFACE effect estimates based on single-varant prediction models. The y-axis denotes the 6 pQTL credible set used to predict TIMP4 protein levels. For each of the six independent credible sets, we use the variant with the highest PIP in a single variant prediction model. We fit a simple linear regression model six times, changing the TIMP4 predicted protein levels each time. The x-axis denotes the gene-to-trait effect estimate and 95% confidence interval. The $I^2$ statistic quantifies the heterogeneity of the independent effect estimates.

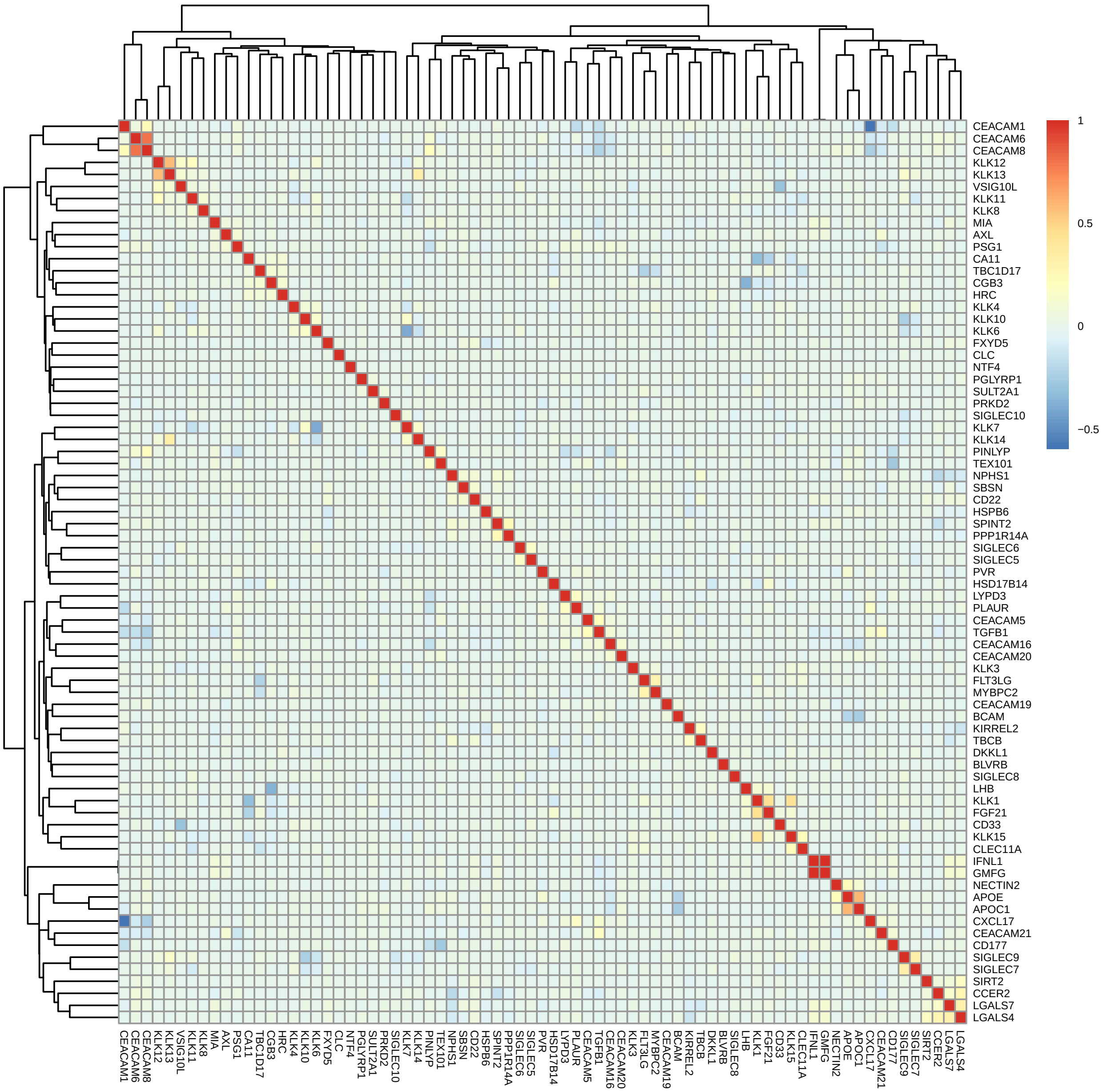


Figure S9: UK Biobank predicted plasma protein levels correlation for all genes proximal to *KLK15*.

### Supplemental Methods

#### INTERFACE prior estimation

This section provides a technical summary for estimating $p_g$, $p_m$, $p_c$, and $\pi$ to construct the INTERFACE priors. Although the estimation procedures described here are not new, they have originated in different application contexts. We include the necessary citations for more detailed references.

**Estimating $p_g, p_m$, and $p_c$** The three quantities are estimated from the enrichment analysis in colocalization analysis. The estimation procedure was first proposed in [1]. It employs a multiple imputation scheme that leverages probabilistic association evidence from the fine-mapping analysis of complex trait GWAS and molecular QTL mapping.

Both $p_g$ and $p_m$ are straightforwardly computed by averaging the SNP PIPs across the genome, i.e.,

$$\hat{p}_g = \frac{\sum_{i=1}^{M} \mathrm{PIP}_{g,i}}{M}$$
$$\hat{p}_m = \frac{\sum_{t=1}^{T} \sum_{k=1}^{K_t} \mathrm{PIP}_{t,k}}{M}$$

where $M$ denotes the total number of SNPs interrogated genome-wide and the estimation for $\hat{p}_m$ is across all $T$ molecular phenotypes, each with $K_t$ *cis*-SNPs. The averaging operation can be interpreted as an application of the law of total expectation, pooling conditional PIPs from all SNPs.

Let the latent binary indicators $d_i$ and $\delta_i$ denote whether SNP $i$ is a causal GWAS hit and a causal molecular QTL, respectively. For instance, $d_i = 1$ and $\delta_i = 1$ indicate that SNP $i$ is a colocalized SNP. To examine the enrichment of molecular QTLs in causal GWAS hits, we consider the following logistic model:

$$\mathrm{logit}\left[\Pr(d_i = 1)\right] = \alpha_0 + \alpha_1 \delta_i, \quad i = 1, 2, \ldots, M$$

Given the values of $\delta_i$'s, the enrichment parameters $(\alpha_0, \alpha_1)$ can be estimated using an EM algorithm, treating $d_i$'s as missing data. In our application, the $\delta_i$'s are unobserved, and only their posterior distributions are available from the QTL fine-mapping analysis. Wen et al. propose using a multiple imputation scheme that directly samples $\delta_i$'s to create multiple datasets with simulated but fixed $\delta_i$ values. Subsequently, the EM algorithm is applied to each simulated dataset to estimate the corresponding enrichment parameters. Finally, the multiple enrichment estimates are combined to produce the final estimate $(\hat{\alpha}_0, \hat{\alpha}_1)$. Note that,

$$\begin{aligned} p_c &= \Pr(d_i = 1 \text{ and } \delta_i = 1) \\ &= \Pr(d_i = 1 \mid \delta_i = 1) \Pr(\delta_i = 1) \\ &= \frac{\exp(\alpha_0 + \alpha_1)}{1 + \exp(\alpha_0 + \alpha_1)} p_m \end{aligned}$$

Thus, we estimate $p_c$ by

$$\hat{p}_c = \frac{\exp(\hat{\alpha}_0 + \hat{\alpha}_1)}{1 + \exp(\hat{\alpha}_0 + \hat{\alpha}_1)} \hat{p}_m$$

**Estimating $\pi$** We apply Storey's procedure to estimate $\pi$ from the TWAS $p$-values. Specifically, we consider the $p$-values from a TWAS analysis as a mixture, where a proportion $\pi$ is generated from the TWAS genes and follows some unknown distribution $G$, and the remaining proportion $1-\pi$ comes from the null genes and follows the uniform distribution. Consider the proportion of the TWAS $p$-values that are no less than 0.5, namely $f_m$. By the law of large number, it follows that

$$f_m \to \frac{1}{2}(1-\pi) + f_{G_m}\pi,$$

where $f_{G_m}$ denotes $P(p \geq 0.5)$ under the $G$ distribution. Without knowing $f_{G_m}$, we note that following inequality should hold when the number of tested genes are sufficiently large,

$$f_m \geq \frac{1}{2}(1-\pi)$$

Thus, we obtain a lower estimate

$$\hat{\pi} = 1 - 2f_m$$

We apply the Storey's procedure to estimate $\pi$ from the TWAS $p$-values. Specifically, we consider the $p$-values from a TWAS analysis as a mixture, where a proportion $\pi$ is generated from the TWAS genes and follows some unknown distribution $G$, while the remaining proportion $1-\pi$ comes from the null genes and follows the uniform distribution. Consider the proportion of the TWAS $p$-values that are at least 0.5, denoted as $f_m$. By the law of large numbers, it follows that

$$f_m \to \frac{1}{2}(1-\pi) + f_{G_m}\pi,$$

where $f_{G_m}$ denotes $P(p \geq 0.5)$ under the $G$ distribution. Without knowing $f_{G_m}$, we note that the following inequality holds when the number of tested genes is sufficiently large:

$$f_m \geq \frac{1}{2}(1-\pi).$$

Thus, we obtain a lower-bound estimate:

$$\hat{\pi} = 1 - 2f_m.$$

#### METSIM Metabolon Metabolite GWAS data

The METSIM study comprises 10,197 men in Kuopio, Finland. Participants aged 45 to 74 were examined in baseline visits from 2005 to 2010. Those who were non-Finnish (n=21), failed whole-genome sequencing (WGS) (n=65), had sex mismatch(n=3), and/or lacked body mass index measurements (n=1) were excluded. Metabolon, Inc (Durham, North Carolina, USA)[2] performed non-targeted metabolomics profiling on EDTA-plasma samples. Samples were obtained after ≥10-hour overnight fasts during baseline visits. First, methanol extraction of biochemicals was applied; then, non-targeted relative quantitative liquid chromatography–tandem mass spectrometry Metabolon DiscoveryHD4 platform was applied to assay 1,544 metabolites. A randomized batch design was used, where batches contained ∼144 METSIM samples and 20 well-characterized human-EDTA plasma samples for quality control. Data processing, including peak quantification and data scaling, was performed for all 10.188 samples together. We used area under the curve to quantify raw mass spectrometry peaks for each metabolite. Overall process variablility was evaluated by the median relative standard deviation for endogenous metabolites that were present in all 20 technical replicates in each batch. In order to adjust for variation caused by day-to-day instrument tuning differences and columns used for biochmical extraction, we scaled the raw peak quantification to the median for each metabolite by batch.

Illumina HiSeq X Ten instruments were used for WGS, targeting a mean depth of at least 30x (paired-end, 150 bp reads). PCR-free library preparation kits from KAPA Biosystems were used for all sequencing. To process samples, CBCL files were converted to FASTQ-formatted reads. Reads were assigned to samples using bcl2fastq conversion software (Illumina Inc., San Diego, CA). Sample-specific FASTQ files were aligned to the GRCh38 genome reference with BWA-mem. Aligned reads were evaluated in BAM files. The Picard MarkDuplicates tool was usied to identify and flag duplicate reads. GVCF files for each individual sample The WeCall variant caller was used to produce GVCF files for each individual sample, which identified SNVs and INDELs as compared to the reference. The samtools tool was used to convert BAM files to CRAM files.

The following criteria were used for quality control: sex discrepancy between genetically-determined and self-reported sex, high rate of heterozygosity/contamination, low sequencing coverage, genetically-identified duplicates, and discordance between whole-exome sequencing and array sequencing.

### Supplemental Results

#### INTACT-INTERFACE difference set PCG-metabolite pairs

In this section, we provide details on some of the gene-metabolite pairs that are implicated by INTACT but INTERFACE and not discussed in the main text. We discuss potential explanations for the discrepancy for most of the cases.

##### VGF-C100001293

The pair VGF-C100001293 has INTERFACE PIP, INTACT posterior, and GRCP of 0.040, 1, and 0.699, respectively. The pair is not validated by the KBA. There are 3 variants with nonzero PIPs in GWAS, pQTL, and INTERFACE fine-mapping. The variant 7:101159880:C:T has a PIP of 1 in GWAS, 0.59 in pQTL analysis, and 1 in INTERFACE. The SNP 7:101173066:C:G has a PIP of 0.16 in GWAS, 0.17 in pQTL analysis, and 0.07 in INTERFACE. The SNP 7:101174620:A:G has a PIP of 0.61 in GWAS, 0.18 in pQTL analysis, and 0.54 in INTERFACE. There are no genes in the model with higher PIP than VGF. We perform a sensitivity analysis similar to the EPHX2 example in the main text using the three independent credible sets in the VGF pQTL fine-mapping results. The heterogeneous effect estimates ($I^2 = 0.997$, Figure S3) provide evidence of pleiotropy.

### CD38-C100001466

The pair CD38-C100001466 has INTERFACE PIP, INTACT posterior, and GRCP of 0.024, 1, and 0.923 respectively. The pair is not validated by the KBA. CD38 is in the same region as BST1, which is implicated by INTERFACE, INTACT, and KBA. Although both CD38 and BST1 have substantial coloclization evidence, the colocalization signals for the two genes come from distinct GWAS fine-mapping credible sets. Interestingly, although there is a modest correlation between predicted protein levels of CD38 and BST1 ($\rho = -0.2$, p-value $< 2.2 \times 10^{-16}$), the PIP for CD38 decreases to $< 0.001$ when we remove BST1 predicted protein levels from the original INTERFACE model. Multi-gene TWAS (i.e., refitting the INTERFACE model without the region genotype matrix) yields PIPs of 0 and 1 for CD38 and BST1, repsectively. This example may represent the linkage [3] between these two paralogues.

**C100004541-AOC1**

The pair C100004541-AOC1 has INTERFACE PIP, INTACT posterior, and GRCP of $< 0.001$, 1, and 0.941, respectively. The pair is validated by the KBA. The variant 7:150857465:C:T has a PIP of 0.79 in GWAS, 1 in pQTL analysis, and 0.69 in INTERFACE. In sensitivity analysis, we find heterogeneous effect estimates ($I^2 = 0.961$, Figure S4), suggesting potential pleiotropy.

**C100010923-APOE**

The pair C100010923-APOE has INTERFACE PIP $< 0.001$, INTACT posterior 0.106, and GRCP 0.9226. The pair is validated by the KBA. APOC1 is in the same region as APOE, which is implicated by INTERFACE, INTACT, and KBA. APOE and APOC1 are tightly linked genes with related biological functions [4]. The colocalization signals for both C100010923-APOE and C100010923-APOC1 come from the same SNP (19:44927023:C:G), and the correlation of the two genes' predicted protein levels is 0.59 (p-value $< 2.2 \times 10^{-16}$, Figure S9). When we refit the INTERFACE model after removing the predicted protein levels of APOC1, the PIP for APOE increases to 0.541. Additionally, multi-gene TWAS (i.e., refitting the INTERFACE model without the region genotype matrix) yields PIPs or 1 for both genes. We observe similar characteristics to those of the C999926054/KLK15/KLK1 case described in the main text; however, in this case, we can confirm that both genes are truly causal for the metabolite. This case suggests that in scenarios where there are two related causal proteins, INTERFACE may be effective at detecting the only one signal at a time.

**C100020487-ADAM8**

The pair C100020487-ADAM8 has INTERFACE PIP $< 0.001$, INTACT posterior 1, and GRCP 0.520. The pair is not validated by the KBA. The variant 10:133339612:A:G is the sole variant in the pQTL credible set responsible for the colocalization signal, but none of the variants prioritized by INTERFACE are members of this credible set. The INTERFACE model prioritized 10:133403671:C:T PIP =0.96, 10:133383872:G:A PIP = 0.55, 10:133383832:A:T PIP= 0.39, and 10:133072424:G:A PIP = 0.30. We have not been able to formulate an explanation for this discrepancy.

##### C100021198-CHMP1A

The pair C100021198-CHMP1A has INTERFACE PIP 0.021, INTACT posterior 1, and GRCP 0.517. The pair is not validated by the KBA. There are 4 variants with nonzero PIPs for pQTL, GWAS, and INTERFACE fine-mapping. 16:89631217:C:T has pQTL PIP 0.04, GWAS PIP 0.11, and INTERFACE PIP 0.19. 16:89639923:A:AC has pQTL PIP 0.32, GWAS PIP 0.23,and INTERFACE PIP 0.13. 16:89641816:T:G has pQTL PIP 0.31, GWAS PIP 0.19, and INTERFACE PIP 0.15. 16:89646481:C:T has pQTL PIP 0.30, GWAS PIP 0.14, and INTERFACE PIP 0.14. These four variants make up the CHMP1A pQTL credible set that contributes to the colocalization signal. In sensitivity analysis, the two CHMP1A pQTL credible sets yield distinct effect size estimates ($I^2 = 0.984$, Figure S5). Due to the lack of pQTL credible sets, it is difficult to assess whether any of the instruments are valid from this analysis. Pleiotropy is a possible explanation for the discrepancy between INTACT and INTERFACE.

##### C1021-OPLAH

The pair C1021-OPLAH has INTERFACE PIP 0.105, INTACT posterior 1, and GRCP 0.833. This pair is validated by the KBA. There are 2 variants with nonzero PIPs for pQTL, GWAS, and INTERFACE fine-mapping. 8:144056712:C:A has pQTL PIP 0.81, GWAS PIP 0.48, and INTERFACE PIP 0.51. 8:144064032:G:A has pQTL PIP 0.19, GWAS PIP 0.51, and INTERFACE PIP 0.49. These two variants make up the OPLAH pQTL credible set that contributes to the colocalization signal. In sensitivity analysis, we find heterogenous effect estimates ($I^2 = 0.970$ Figure S5), suggesting potential pleiotropy.

##### C999918899-EPHX2

The pair C999918899-EPHX2 has INTERFACE PIP 0.251, INTACT posterior 0.225, and GRCP 0.603. This pair is not validated by the KBA. There are 3 variants with nonzero PIPs for pQTL, GWAS, and INTERFACE fine-mapping. 8:27495233:A:G has pQTL PIP 0.02, GWAS PIP 0.06, and INTER-

FACE PIP 0.04. 8:27516348:G:A has pQTL PIP 0.65, GWAS PIP 0.85, and INTERFACE PIP 0.50. 8:27517169:A:T has pQTL PIP 0.07, GWAS PIP 0.07, and INTERFACE PIP 0.05. These variants are contained within a single EPHX2 pQTL credible set and drive the colocalization signal. In sensitivity analysis, we find heterogenous effect estimates ($I^2 = 0.934$, Figure S7), suggesting potential pleiotropy.

##### C999924328-TIMP4

The pair C999924328-TIMP4 has INTERFACE PIP 0.002, INTACT posterior 1, and GRCP 0.794. This pair is not validated by the KBA. There are 2 variants with nonzero PIPs for pQTL, GWAS, and INTERFACE fine-mapping. 3:12114388:T:C has pQTL 0.82, GWAS PIP 0.4 , and INTERFACE PIP 0.41. 3:12148468:A:G has pQTL PIP 0.17, GWAS PIP 0.6, and INTERFACE PIP 0.59. These variants represent a single TIMP4 credible set responsible for the colocalization signal. In sensitivity analysis, we find heterogenous effect estimates ($I^2 = 0.915$, Figure S8), suggesting potential pleiotropy.
